# Krüppel-like factor 4 is required for development and regeneration of germline and yolk cells from somatic stem cells in planarians

**DOI:** 10.1101/2021.11.08.467675

**Authors:** Melanie Issigonis, Akshada B. Redkar, Tania Rozario, Umair W. Khan, Rosa Mejia-Sanchez, Sylvain W. Lapan, Peter W. Reddien, Phillip A. Newmark

## Abstract

Sexually reproducing animals segregate their germline from their soma. In addition to gamete-producing gonads, planarian and parasitic flatworm reproduction relies on yolk-cell-generating accessory reproductive organs (vitellaria) supporting development of yolkless oocytes. Despite the importance of vitellaria for flatworm reproduction (and parasite transmission), little is known about this unique evolutionary innovation. Here we examine reproductive system development in the planarian *Schmidtea mediterranea,* in which pluripotent stem cells generate both somatic and germ cell lineages. We show that a homolog of the pluripotency factor Klf4 is expressed in primordial germ cells, presumptive germline stem cells, and yolk-cell progenitors. *klf4* knockdown animals fail to specify or maintain germ cells; surprisingly, they also fail to maintain yolk cells. We find that yolk cells display germ-cell-like attributes and that vitellaria are structurally analogous to gonads. In addition to identifying a new proliferative cell population in planarians (yolk cell progenitors) and defining its niche, our work provides evidence supporting the hypothesis that flatworm germ cells and yolk cells share a common evolutionary origin.

## Introduction

Sexually reproducing animals consist of two main cell types: germ cells that produce gametes (eggs and sperm), and somatic cells that make up the remainder of the body. Animal germ cells are typically specified in either of two ways: by determinate or inductive specification. Determinate specification results from the segregation of specialized maternal determinants (germ plasm) at the onset of embryogenesis; those cells receiving germ plasm acquire germ cell fate. In contrast, inductive specification occurs later in embryogenesis when extrinsic signals from surrounding tissues instruct competent cells to form germ cells. Determinate specification has been studied extensively in traditional laboratory models, including *Drosophila*, *C.elegans*, zebrafish, and frogs [1–4]. Inductive specification is less well characterized, even though it is the basal and most common mode of germ cell specification in the animal kingdom, and it occurs in mammals [2, 3].

Irrespective of the mode of germ cell specification, an important commonality exists: once formed, germ cells are set aside from the soma. The developmental decision to segregate the germ cell lineage from somatic cells is essential for species continuity; unlike the soma, which expires each generation, “immortal” germ cells pass on genetic information and serve as a perpetual link between generations. Many animals (e.g., *Drosophila*, *C. elegans*, and mice) specify their germ cells (and segregate them from their soma) only once during embryonic development [1–4]. However, some animals retain the ability to specify new germ cells throughout their lifetime. Sponges and cnidarians maintain into adulthood multipotent stem cells that fuel the continuous production of new germ cells while also giving rise to somatic cell lineages [5–11]. How do these stem cells decide between somatic and germ cell fates?

Planarian flatworms can regenerate an entire body from small tissue fragments. Intensive efforts have been devoted to understanding the mechanisms underlying this regenerative prowess. Planarian regeneration is driven by pluripotent stem cells called neoblasts that are distributed throughout the body [12–14]. Planarians can also inform our understanding of germ cell biology: the neoblasts that give rise to all somatic lineages also give rise to new germ cells. Interestingly, neoblasts and germ cells express a shared set of conserved “germline genes,” including *piwi*, *vasa*, *pumilio*, and *tudor* [15, 16], which play important roles in neoblast function [17–27]. Like mammals, planarians undergo inductive germ cell specification [28–30]. However, the mechanistic basis underlying germ cell specification from “somatic” neoblasts and the factors involved in adopting somatic versus germ cell fate remain obscure.

Here we investigate how new germ cells are specified from neoblasts throughout post-embryonic development and during regeneration in planarians. We also examine another critical aspect of the planarian reproductive system: the development of an extensive network of accessory organs known as vitellaria. Unique among animals, eggs of most flatworms are ectolecithal: yolk is not present within oocytes themselves, but rather is made by vitellaria that produce specialized yolk cells (vitelline cells or vitellocytes). Planarians and all parasitic flatworms are characterized by ectolecithality. However, despite the importance of vitellaria in the life cycle and transmission of these parasites [31, 32], little is known about the development of vitellaria or production of yolk cells.

We show that a homolog of the conserved transcription factor *krüppel-like factor 4* (*klf4*), a critical inducer of pluripotency in mammals [33], is expressed in male and female presumptive germline stem cells (GSCs) in the planarian *Schmidtea mediterranea*, as well as in a newly discovered population of mitotically competent yolk cell progenitors. We demonstrate that *klf4* is required for germ cell specification and that *klf4* knockdown leads to the loss of both germ cell and yolk cell lineages. We provide evidence that yolk-cell-producing organs in planarians consist of two distinct cell types: a yolk cell lineage, which is characterized by several germ-cell-like attributes, and support cells, which sustain yolk cell maintenance and differentiation. Our data show that planarian vitellaria are structurally analogous to gonads and that yolk cells share several important features with both somatic neoblasts and germ cells.

## Results

### *klf4* is expressed in planarian gonads and yolk-producing accessory organs

In the search for regulators of germ cell fate in planarians, we focused on the conserved transcription factor KLF4, a key pluripotency factor in mammals [33]. Sexual *S. mediterranea* are cross-fertilizing, simultaneous hermaphrodites. Using fluorescent in situ hybridization (FISH) on sexually mature adults, we found that *klf4* mRNA is expressed at high levels within the ventrally situated ovaries, as well as in cells that are distributed along the medial-posterior region of each lobe of the cephalic ganglia and appear to be arranged in a field anterior to each ovary. Sparse *klf4^+^* cells are also located dorsolaterally, where the testes reside (Fig 1A and 1B). Additionally, *klf4^+^* cells are scattered throughout the ventral parenchyma (the tissue surrounding the planarian’s internal organs), in a pattern reminiscent of vitellaria, the yolk-producing organs essential for reproduction (Fig 1A and 1B). Thus, this pluripotency-associated transcription factor is expressed in male and female reproductive tissues.

**Fig 1.**
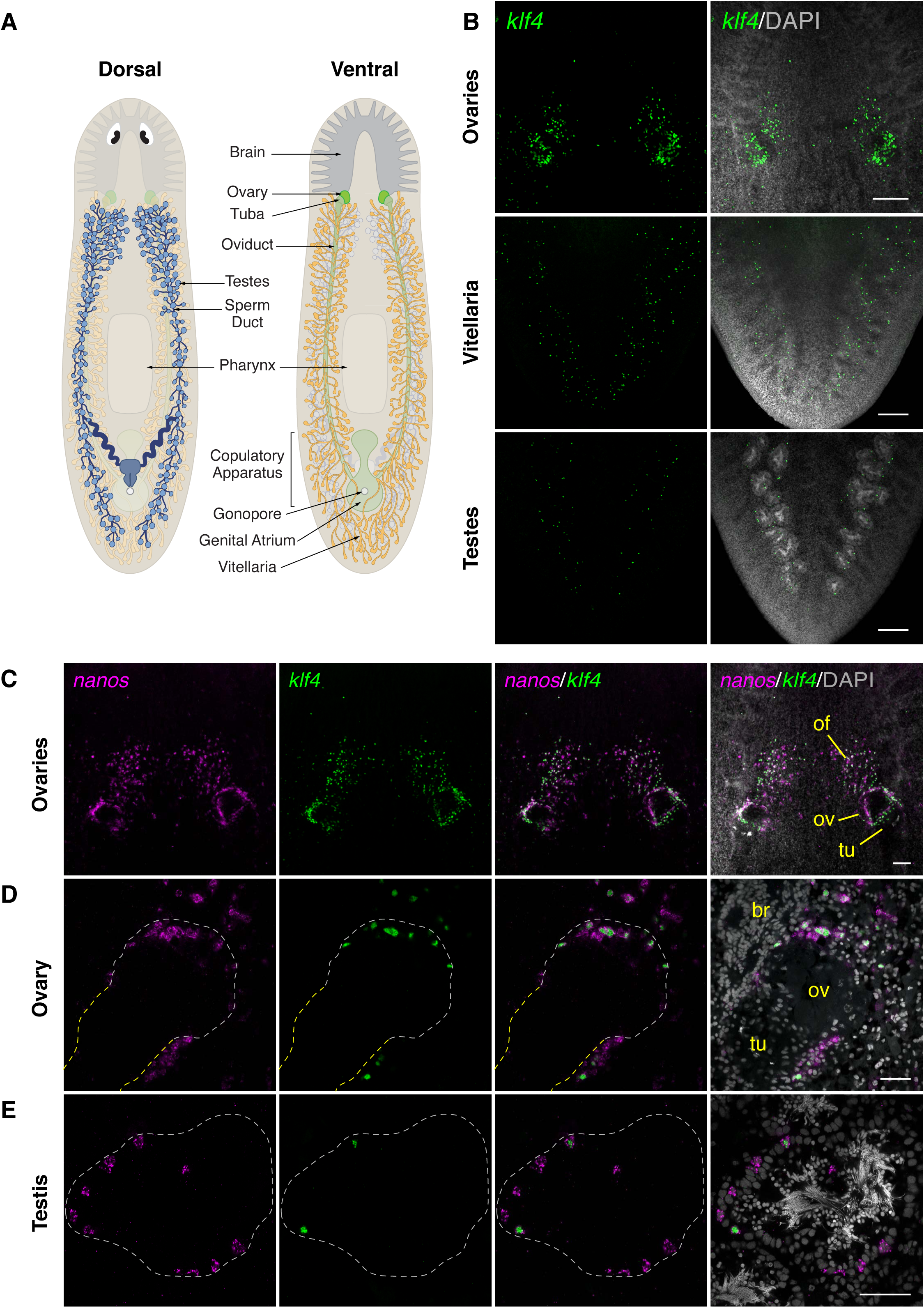
*klf4* is expressed in gonads and vitellaria and is restricted to a subset on *nanos^+^* germ cells in planarian ovaries and testes. (A) Schematics depicting the dorsal (left) and ventral (right) views of landmark structures and various reproductive organs in adult sexual *S. mediterranea*. (B) Maximum-intensity projections of confocal sections showing fluorescence in situ hybridization (FISH) of *klf4* (green) in ventral head region (top), ventral tail region (middle), and dorsal tail region (bottom). (C) Maximum-intensity projection of confocal sections showing double FISH (dFISH) of *klf4* (green) and germline marker *nanos* (magenta) in ventral head region. *klf4-* and *nanos*-expressing cells are detected surrounding the tuba (tu) at the base of each ovary (ov), along the periphery of the ovaries, and in anterior ovarian fields (of) situated mediolaterally along the brain. (D) Single confocal section of a planarian ovary located posterior to the brain (br) showing *klf4* (green) and *nanos* (magenta) dFISH. *klf4-* and *nanos*-expressing cells are found at the ovary-tuba junction, along the periphery of the ovary, and in germ cells anterior to the ovary. Dashed line denotes ovary (white) and tuba (yellow) boundary. (E) Confocal section of *klf4* (green) and *nanos* (magenta) dFISH showing *klf4*/*nanos* double-positive and *nanos* single-positive cells along the periphery of the testis. Dashed line denotes testis boundary. (B-E) Nuclei are counterstained with DAPI (gray). Scale bars, 200 µm (B), 100 µm (C), and 50 µm (D-E).

### *klf4* expression is restricted to a subset of *nanos^+^* germ cells in ovaries and testes

To analyze gonadal *klf4* expression in more detail, we performed double FISH (dFISH) to detect *klf4* and the germline marker *nanos* [34] (Fig 1C-1E). Previous work in planarians has shown that gonadal *nanos* expression is restricted to the early spermatogonia and oogonia in the outermost layer of testes and ovaries, respectively, which have been interpreted as presumptive germline stem cells (GSCs) [29,35–37]. In addition to the previously described *nanos^+^* germ cells found at the ovary periphery, we detected *nanos^+^* cells in the same anterior ovarian fields described above (Fig 1C); a substantial proportion of *nanos^+^* cells in these fields co-expresses *klf4* (81%, n=1116 *nanos^+^* cells) and all *klf4^+^* cells are *nanos^+^* (100%, n=908 *klf4^+^* cells). *klf4* expression is similarly restricted to a subset of *nanos^+^* germ cells located at the ovary periphery, and in *nanos^+^* cells clustered at the boundary between the ovary and its outlet, the tuba (the anterior-most portion of the oviduct where fertilization occurs) (Fig 1D) (90%, n=1588 *nanos^+^* cells). All *klf4^+^* cells within and at the base of the ovary co-express *nanos* (100%, n=1423 *klf4^+^* cells). In the testes of sexually mature adults, *klf4* is also expressed around the periphery, but is confined to a small subset of *nanos^+^* germ cells (14%, n=10,628 *nanos^+^* cells) (Fig 1E). Similar to female germ cells, all *klf4^+^* male germ cells express *nanos* (100%, n=1475 *klf4^+^* cells). Our observations show that in both ovaries and testes, the *nanos^+^* presumptive GSCs are heterogeneous with respect to *klf4* expression.

Since only a fraction of *nanos^+^* germ cells express *klf4*, we wondered whether *klf4* expression represents the earliest stages of *nanos^+^* germ cell development. To answer this question, we examined the developmental progression of *klf4* and *nanos* expression, starting from the emergence of primordial germ cells (PGCs) in newly hatched planarians. Previous studies describing *nanos* expression in hatchlings failed to detect the presence of female (i.e., anteroventral) PGCs in planarians until 1 week post-hatching. Male (dorsolateral) *nanos^+^* PGCs were observed in a minority of planarians during the final stages of embryonic development (stage 8 embryos) and in 1-day-old hatchlings [34, 36]. In contrast to these studies, by FISH, we were able to detect female *nanos^+^* cells in 100% of 1-day-old hatchlings; however, only a fraction of these cells expresses the neoblast/germline marker *piwi-1*, (40%, n=1199 *nanos^+^* cells), indicating that not all anteroventral *nanos^+^* cells are germ cells. While predominantly expressed in germ cells, *nanos* transcripts have also been detected in a population of eye cells [36]. Consistent with their ventral location and proximity to the cephalic ganglia, we postulate that *nanos^+^*/*piwi-1^−^* cells may represent another somatic cell population, such as neurons. To determine the proportion of *nanos^+^* PGCs that express *klf4*, we performed triple FISH and found that 56% of *nanos^+^*/*piwi-1^+^* PGCs co-express *klf4* (n=598 *nanos^+^*/*piwi-1^+^* cells) (S1A Fig), indicating that this heterogeneity persists throughout sexual development (72% and 75% of *nanos^+^*/*piwi-1^+^* germ cells are *klf4^+^* in immature, juvenile ovaries and mature, adult ovaries, respectively) (S1B Fig).

Male germ cells are easily observed throughout testis maturation. In hatchlings, all *nanos^+^* cells distributed dorsolaterally (where testes will develop) co-express *piwi-1* (100%, n=517 *nanos^+^* PGCs). Essentially all *nanos^+^* PGCs also co-express *klf4* (98%, n=189 *nanos^+^* cells) (Fig 2A). However, as planarians undergo sexual maturation and testis primordia continue to develop, *klf4* is expressed in an increasingly smaller proportion of the *nanos*^+^ population (34%, n=5198 *nanos^+^* cells in juveniles and 14%, n=10,628 *nanos^+^* cells in adults) and there is a marked increase in *klf4^−^*/*nanos^+^* germ cells (Fig 2B-2C). These data indicate that in hatchlings, newly specified PGCs express both *klf4* and *nanos*; whereas, during sexual development a *nanos* single-positive germ cell population emerges and expands. These observations are consistent with a model in which *klf4^+^*/*nanos^+^* cells represent the most undifferentiated germ cell state (i.e. PGC and GSC), and *klf4^−^*/*nanos^+^* germ cells are their immediate progeny.

**Fig 2.**
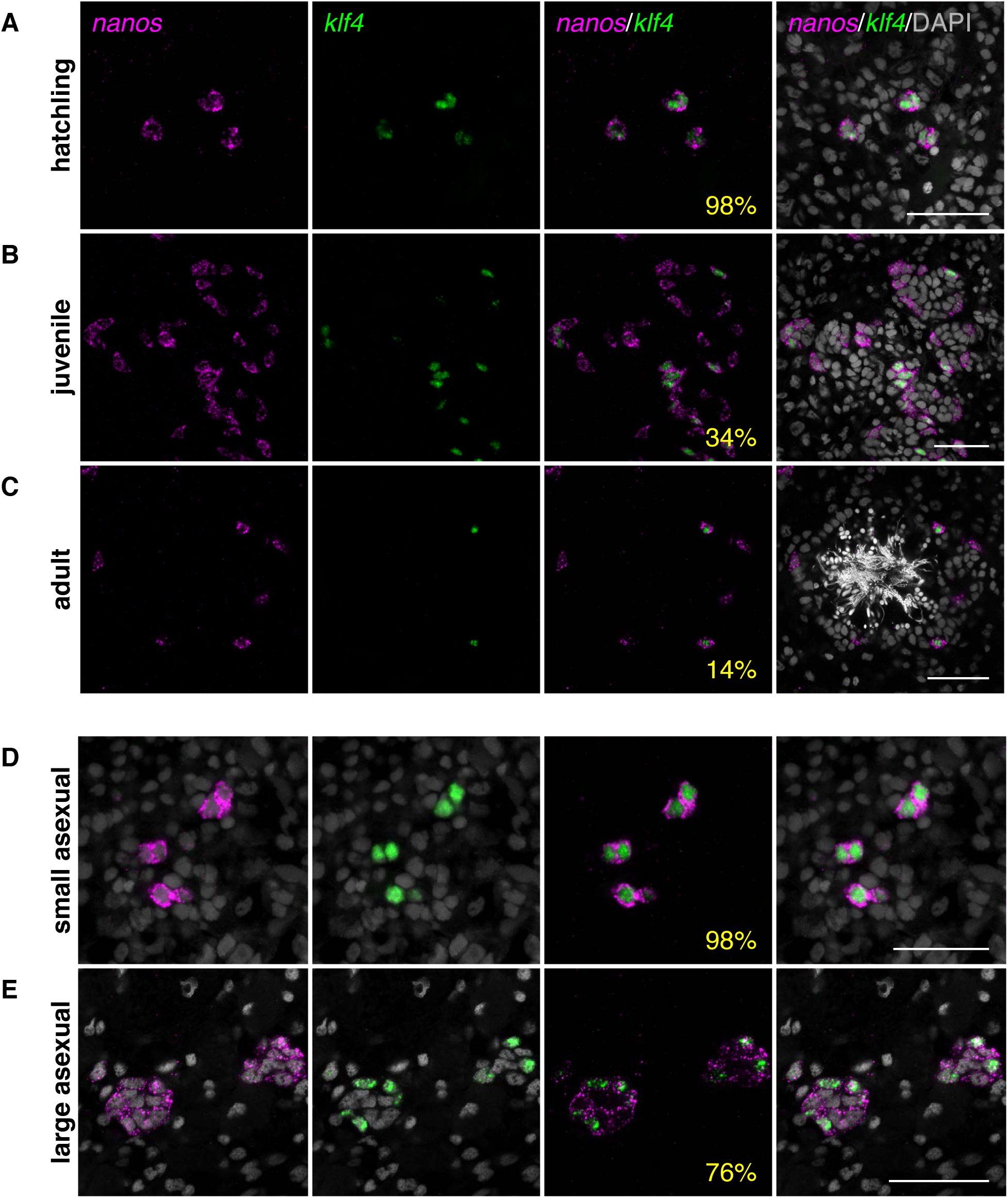
*klf4* expression becomes restricted to a subset of *nanos^+^* male germ cells during sexual planarian maturation and asexual planarian growth. (**A-C**) Confocal sections showing dFISH of *klf4* (green) and *nanos* (magenta) in hatchling testis primordia (**A**), juvenile testes (**B**), and sexually mature adult testes (**C**). *klf4* is expressed in most *nanos^+^* male PGCs in hatchlings and becomes progressively restricted to a subpopulation of *nanos^+^* germ cells as planarians sexually mature. (**D-E**) Confocal sections showing dFISH of *klf4* (green) and *nanos* (magenta) in testis primordia in small (**D**) and large (**E**) asexual planarians. *klf4* is co-expressed in almost all *nanos^+^* male germ cells in small asexuals and is restricted to a subset of *nanos^+^* male germ cells in large asexuals. (**A-E**) Percentages reflect *nanos^+^* cells that are also *klf4^+^*. Nuclei are counterstained with DAPI (gray). Scale bars, 50 µm.

Thus far, we have characterized *klf4* expression in the sexual strain of *S. mediterranea*. However, this species also exists as an obligate asexual biotype, which reproduces exclusively by fission and does not produce mature gametes or accessory reproductive organs. Although asexual planarians do not develop functional gametes, they nevertheless specify PGCs in small clusters of *nanos^+^* gonadal primordia. These *nanos^+^* cells do not proliferate or differentiate [34,36,37]. By comparing small (∼2 mm) and large (>5 mm) asexuals, we found that the number of *nanos^+^* germ cells in female (anteroventrally located) and male (dorsolaterally located) primordia increases as animals grow (S1C and S1D Fig). We examined whether *klf4* expression was restricted to these early PGCs, and by dFISH we found that *klf4* is co-expressed in the majority of female *nanos^+^* cells, in similar proportions for both small and large asexuals (91%, n=24 *nanos^+^* cells and 86%, n=213 *nanos^+^* cells, respectively) (S1C and S1D Fig). In contrast, *klf4* is co-expressed in virtually all male *nanos^+^* cells in small asexuals (98%, n=559 *nanos^+^* cells), whereas testis primordia in larger animals contain both *klf4^+^*/*nanos^+^* cells and *klf4^−^*/*nanos^+^* cells (76% and 24% respectively, n=1645 *nanos^+^* cells) (Fig 2D and 2E). Thus, in both growing asexuals and maturing sexuals, as *nanos^+^* cells in testis primordia increase in number, *klf4* expression becomes restricted to a subset of these germ cells. This similarity suggests that in the asexual biotype germ cells can undergo the first step of development – from *klf4^+^*/*nanos^+^* to *klf4^−^*/*nanos^+^* cells – before reaching a block in differentiation.

### *klf4*-expressing germ cells in ovaries and testes are mitotically active

In many animals, the production of gametes in adulthood is enabled by GSCs. Our findings raise the possibility that *klf4*-expressing cells are GSCs representing the top of oogonial and spermatogonial lineages. All GSCs have the ability to undergo self-renewing divisions, which give rise to differentiating daughter cells while maintaining the stem cell pool. By combining phospho-Histone H3 (pHH3) immunostaining with *klf4* and *nanos* dFISH, we examined the mitotic profiles of cells within the germ cell hierarchy and sought to ascertain whether *klf4^+^*/*nanos^+^* cells are competent to divide and, therefore, fulfill a basic criterion of GSC behavior.

We found that *klf4^+^*/*nanos^+^* germ cells within the ovarian fields are mitotically active (0.3%, n=3409 *klf4^+^*/*nanos^+^* cells) (Fig 3A). We also detected proliferation of *klf4^−^*/*nanos^+^* oogonia at the outer periphery of the ovaries, whereas *nanos^−^* oogonia within the ovaries do not divide mitotically (Fig 3B). Thus, female germ cells are specified and proliferate within the ovarian field and/or the ovary periphery, and as oogonia turn off *nanos* expression, they cease to divide mitotically and differentiate into oocytes.

**Fig 3.**
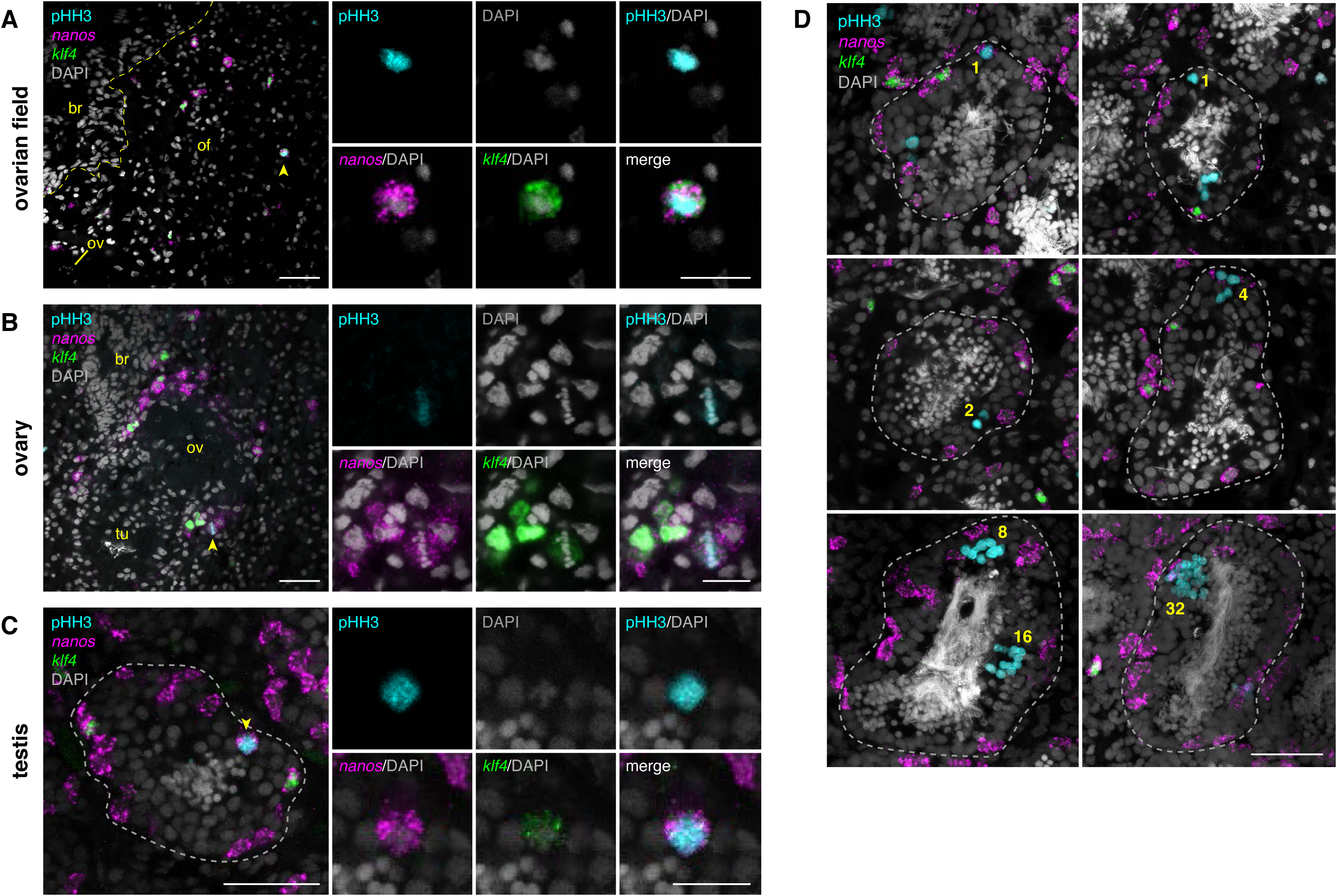
*klf4^+^* germ cells in planarian ovaries and testes are mitotically active. (**A-C**) Confocal sections showing dFISH of *klf4* (green) and *nanos* (magenta) and immunostaining of mitotic marker phospho-Histone H3 (pHH3; cyan) in the ovarian field (of) located anterior to the ovary (ov) and proximal to the brain (br; boundary denoted by yellow dashed line) (**A**), the ovary, which is anterior to the tuba (tu) (**B**), and the testis (boundary denoted by gray dashed line) (**C**). Side panels are high magnification views of *klf4*/*nanos*/pHH3 triple-positive cells (yellow arrowheads). (**D**) Confocal sections (top 4 panels) and maximum-intensity projections (bottom 2 panels) showing dFISH of *klf4* (green) and *nanos* (magenta) and immunostaining of pHH3 (cyan) in testes. Yellow numbers denote pHH3^+^ germ cells dividing throughout spermatogenesis: single *nanos^+^* cell; single *nanos^−^* cell; 2-, 4-, 8-cell spermatogonial cysts; and 16- and 32-cell spermatocyte cysts. (**A-D**) Nuclei are counterstained with DAPI (gray). Scale bars, 50 µm for whole-gonad images, 20 µm for side panels.

Male germ cells actively divide throughout spermatogenesis; spermatogonia undergo 3 rounds of synchronous mitotic divisions with incomplete cytokinesis to produce 2-, 4-, and 8-cell spermatogonial cysts connected by intercellular bridges, whose cells differentiate into primary spermatocytes and divide meiotically to generate 32 spermatids that ultimately transform into mature sperm [28, 38]. We detected pHH3*^+^*/*klf4^+^*/*nanos^+^* triple-positive cells in testes of both hatchlings (1%, n=773 *klf4^+^*/*nanos^+^* cells) and adults (0.2%, n=2436 *klf4^+^*/*nanos^+^* cells). In mature sexuals, mitotic, single-cell spermatogonia and mitotic doublets were observed in *nanos^+^* germ cells (including *klf4^+^*/*nanos^+^* cells) along the outermost periphery of the testis (Fig 3C and 3D). We also observed *nanos^−^*/pHH3^+^ singlets and doublets, which might represent mitotic *nanos^−^* single-cells or 2-cell spermatogonia (Fig 3D). We never detected *nanos* expression in pHH3^+^ 4- or 8-cell premeiotic spermatogonial cysts, or in 16- or 32-cell meiotic cysts (Fig 3D). All our observations thus far support a model in which the spermatogonial lineage consists of *klf4^+^*/*nanos^+^* germ cells at the top of the hierarchy giving rise to *klf4^−^*/*nanos^+^* and subsequently *klf4^−^*/*nanos^−^* single-cell spermatogonia, and that germ cells cease expressing *nanos* once spermatogonial cystogenic divisions have occurred (Fig 3D).

### *klf4* is required for female and male gametogenesis and is necessary for PGC specification

Having established *klf4* as the earliest germ cell marker, we asked whether *klf4* is required for gonadal development. We induced *klf4* RNAi by feeding hatchlings double-stranded RNA (dsRNA) twice a week for 4-6 weeks – the time normally required to reach sexual maturity. We examined the effects of *klf4* knockdown on gonadal development by FISH to detect markers of early germ cells (*nanos*), oocytes (*Cytoplasmic Polyadenylation Element Binding Protein 1*, *CPEB1*), and gonadal somatic support cells (*ophis*, *Laminin A*, and *dmd1*) (Fig 4A) [35,39,40]. *klf4* knockdown resulted in a significant reduction of early (*nanos^+^*) germ cells in the anterior ovarian fields and ovaries as well as a loss of mature (*CPEB1^+^*) oocytes (Fig 4B and 4C, S2 Fig). In extreme cases, ovaries were devoid of mature germ cells. Despite this dramatic loss of germ cells, *klf4* RNAi ovaries were larger than their control RNAi counterparts, because of a significant expansion of (*ophis^+^* or *LamA^+^*) somatic support cells (Fig 4B and 4C). *klf4* RNAi also led to a loss of germ cells in the testes (Fig 4D and 4E). Agametic *klf4* RNAi testes consist of *dmd1^+^* somatic cells and have a “collapsed” appearance due to the absence of germ cells (Fig 4D and 4E). Therefore, these data indicate that *klf4* is required for the maintenance of *nanos^+^* germ cells to sustain proper gametogenesis in both ovaries and testes.

**Fig 4.**
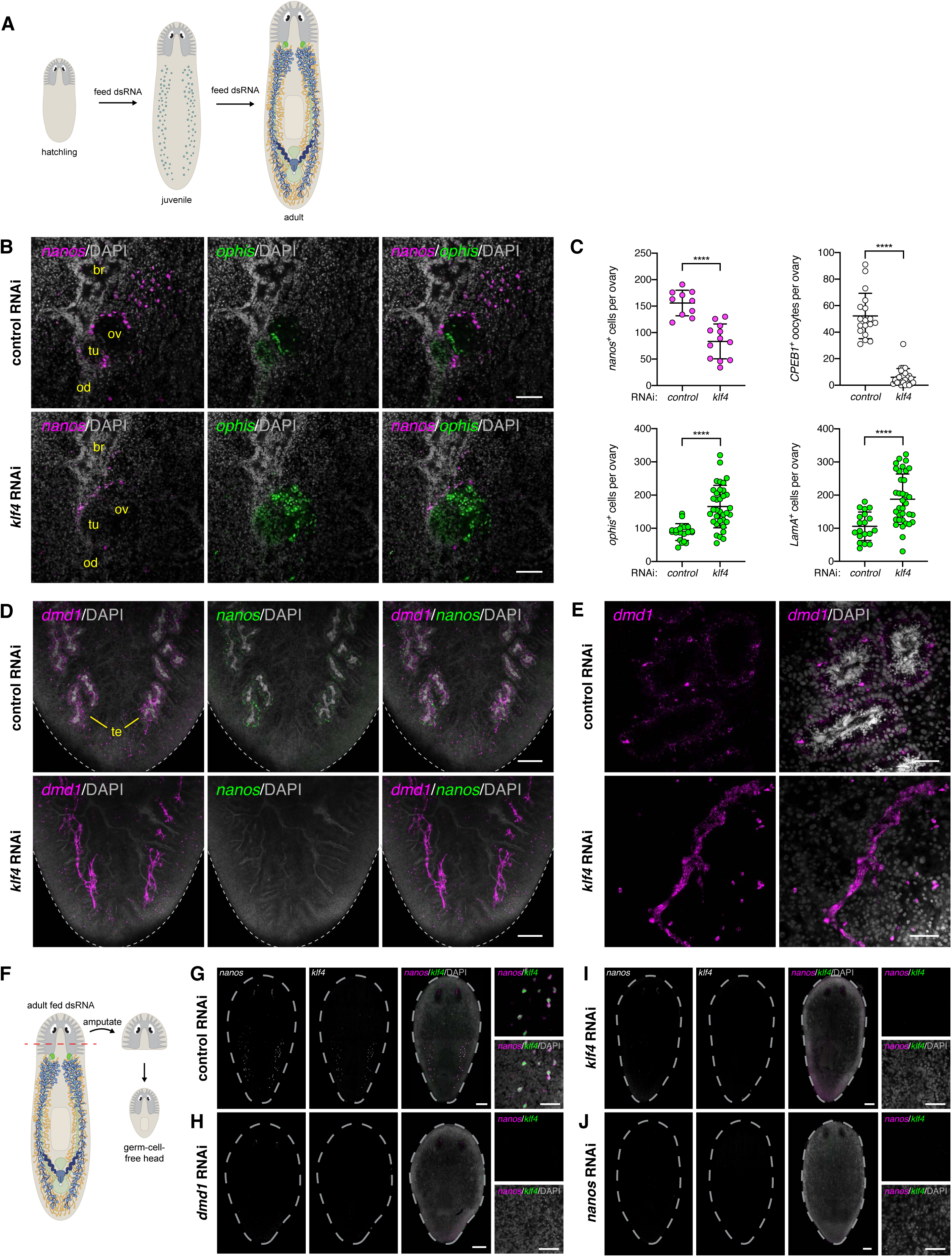
*klf4* is required for gametogenesis in adult ovaries and testes and is necessary for PGC specification. (**A**) RNAi scheme during development in sexual *S. mediterranea* from newborn hatchling to sexually mature adult. (**B**) Single confocal section of an ovary (ov) located at the posterior of the brain (br) showing dFISH of *nanos* (magenta) and *ophis* (green; somatic gonadal cells, tuba (tu), oviduct (od) in control and *klf4* RNAi planarians. (**C**) Quantification of *nanos^+^* germ cells, *CPEB1^+^* oocytes, *ophis^+^* somatic gonadal cells, and *LamA^+^* somatic gonadal cells per ovary in control and *klf4* RNAi animals. Data are presented as mean ± SD. *klf4* RNAi results in significantly fewer germ cells and a corresponding increase in somatic support cells compared to control RNAi ovaries, p<0.0001, two-tailed Welch’s t-test. (**D**) Maximum-intensity projections of confocal sections showing dFISH of *dmd1* (magenta; somatic gonadal cells) and *nanos* (green) in dorsal tail region. *dmd1-* and *nanos*-expressing cells are detected surrounding the DAPI-rich sperm located at the center of each testis (te) in control RNAi planarians. *dmd1^+^* somatic gonadal cells are present but display a “collapsed” appearance due to the loss of germ cells in *klf4* RNAi planarians. Dashed line denotes planarian boundary. (**E**) Confocal sections of control and *klf4* RNAi testes. Note loss of spermatogenesis and “collapsed” appearance of *dmd1^+^* somatic gonadal cells in *klf4* RNAi testes compared to controls. (**F**) Amputation scheme to assay de novo re-specification of germ cells. Amputation anterior to the ovaries results in a head fragment lacking any reproductive tissues (soma only). This head fragment will regenerate a new trunk and tail and will specify new germ cells. (**G-J**) Maximum-intensity projections of confocal sections showing dFISH of *klf4* (green) and *nanos* (magenta) in head regenerates 2 weeks post-amputation. N=3-5 experiments, n=10-35 planarians (**G**) Control RNAi head regenerates specify new *nanos^+^* PGCs that co-express *klf4*. (**H-J**) *klf4* and *nanos* RNAi head regenerates phenocopy *dmd1* knockdowns and fail to specify *klf4^+^*/*nanos^+^* PGCs. (**B, D-E, G-J**) Nuclei are counterstained with DAPI (gray). Scale bars, 100 µm (**B**), 200 µm (**D**), 50 µm (**E**), 200 µm for whole-planarian images, 50 µm for side panels (**G-J**).

Is *klf4* also necessary for PGC specification? Since *klf4^+^*/*nanos^+^* PGCs are specified during embryogenesis and are already present in newborn hatchlings, it is not feasible to induce RNAi by dsRNA feeding before PGC specification. However, planarians can regenerate germ cells de novo – amputated head fragments comprised solely of somatic cells can inductively specify new germ cells [34, 41]. Therefore, to test the requirement of a gene during PGC specification, one can feed adult planarians dsRNA to induce RNAi, amputate heads anterior to all reproductive tissues, and examine regenerating “germ cell-free” head fragments for de novo germ cell specification (Fig 4F) [35]. Two weeks post-amputation, we found that control head fragments re-specified *klf4^+^/nanos^+^* cells dorsolaterally (Fig 4G) [34]. By contrast, re-specification of PGCs in *klf4* RNAi or *nanos* RNAi head fragments was significantly impaired, with several head regenerates lacking germ cells entirely (Fig 4I and 4J). Similarly, *dmd1* RNAi head fragments fail to specify new *klf4^+^*/*nanos^+^* germ cells during regeneration (Fig 4H), because *dmd1* is required non-autonomously for germ cell specification [35]. These data indicate that *klf4* and *nanos* are required cell autonomously for specification of new germ cells from neoblasts and that *klf4^+^*/*nanos^+^* male PGCs rely on somatic gonadal “niche” cells for their induction.

### *klf4*-expressing cells are present in vitellaria and are progenitors of the yolk cell lineage

Our results indicate that *klf4* is an essential regulator of the establishment and maintenance of the planarian germ cell lineage. However, this crucial germline regulator is also expressed in “somatic” organs: the vitellaria (Fig 1B). In *S. mediterranea*, the vitellaria are located ventrally beneath the testes and connect to the oviducts (Fig 1A). Yolkless oocytes are fertilized in the anterior-most compartment of the oviduct (the tuba). After fertilization, zygotes are transported posteriorly through the oviducts to the genital atrium, accumulating thousands of yolk cells along the way. One or more zygotes and numerous extraembryonic yolk cells are then enclosed within egg capsules. These capsules are laid through the gonopore, and embryonic development proceeds for two weeks before newborn hatchlings emerge [28,42–44]. Planarian embryos rely on vitellaria-derived yolk cells for their nutritional needs and development. However, little is known about these essential reproductive structures, or how yolk cells are made.

To our surprise, *klf4*-expressing cells in the vitellaria also expressed *nanos* (97%, n=1548 *klf4^+^* cells) (Fig 5A and 5A’). Previously, *nanos* expression had only been detected in a population of eye cells and in germ cells in testes and ovaries [34,36,37]. Compared to germ cells, *nanos* is expressed at lower levels in the vitellaria, but is readily detectable due to recent improvements in ISH sensitivity [45, 46]. As in the gonads, *klf4* expression in the vitellaria is restricted to a subset of *nanos^+^* cells (49%, n=3304 *nanos^+^* cells). Do *klf4^+^*/*nanos^+^* cells represent the progenitor of planarian yolk cells? To answer this question and to characterize the progression of yolk cell development, we performed combinatorial dFISH analyses on mature sexual planarians to detect *klf4* and previously reported vitellaria markers *CPEB1*, *surfactant b*, and *Monosiga brevicollis MX1 hypothetical protein* (*MX1*) [40]. We found that some *klf4^+^* cells co-express *CPEB1* (10%, n=984 *klf4^+^* cells) and *surfactant b* (9%, n=822 *klf4^+^* cells), but not *MX1* (0%, n=1094 *klf4^+^* cells) (Fig 5B–5D and 5B’–5D’). These observations suggest that *klf4* expression marks the earliest yolk cells and that *CPEB1* and *surfactant b* expression precede *MX1* expression in the yolk cell lineage. Indeed, most *CPEB1^+^* cells co-express *surfactant b* (95%, n=1752 *CPEB1^+^* cells) but a much smaller fraction co-express *MX1* (36%, n=4334 *CPEB1^+^* cells) (S3A, S3B, S3A’ and S3B’ Fig). A large fraction of *surfactant b*-expressing cells are also *MX1^+^* (70%, n=8057 *surfactant b^+^* cells), and essentially all *MX1^+^* cells co-express *surfactant b* (99 %, n=5840 *MX1^+^* cells) (S3C and S3C’ Fig). The progression of yolk cell development is also marked by changes in cell morphology: *klf4^+^* cells are small and have very little cytoplasm, unlike the large, yolk-filled *surfactant b^+^* and *MX1^+^* cells of the vitellaria (Fig 5C’ and 5D’).

**Fig 5.**
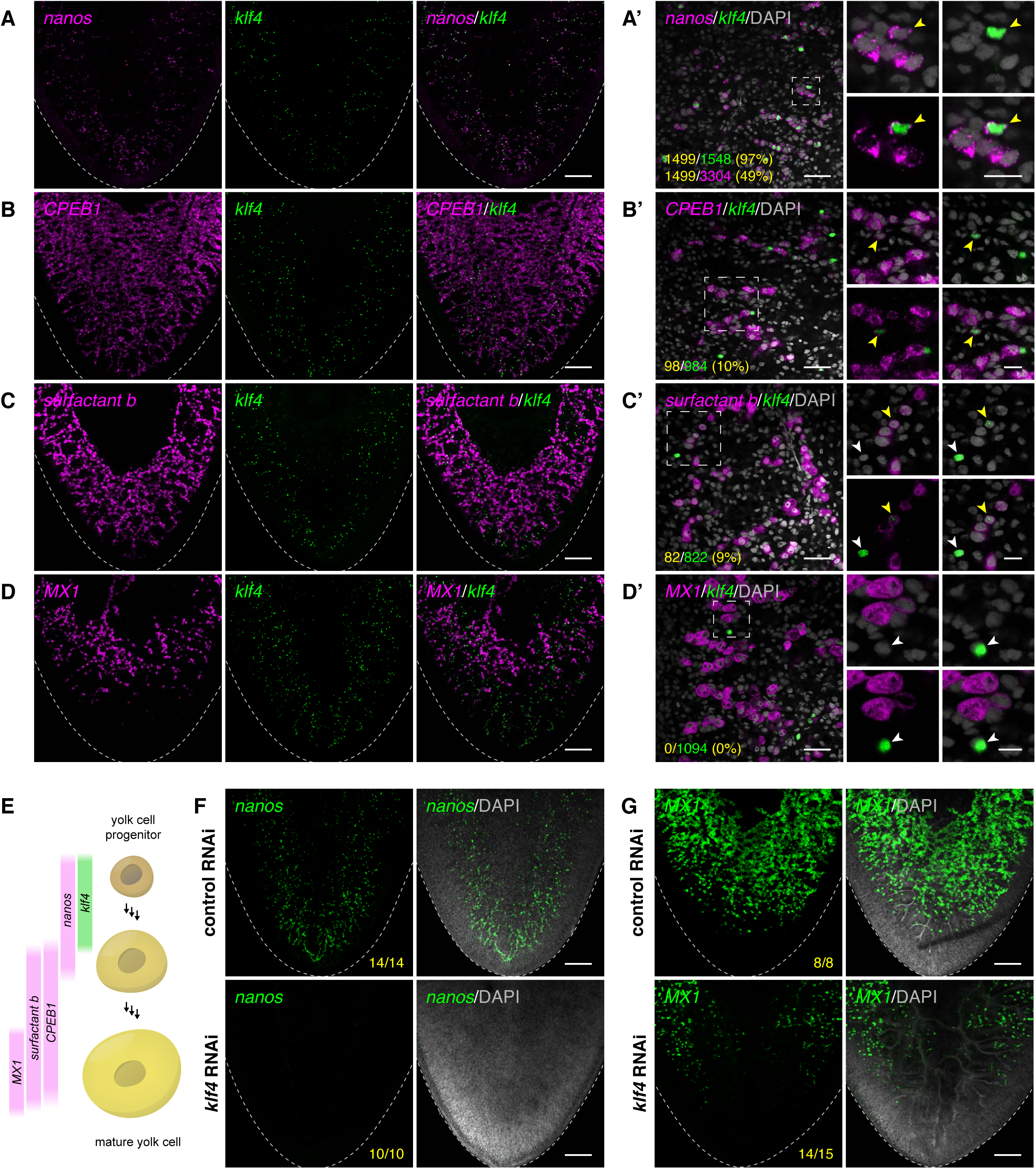
*klf4^+^* cells are present in vitellaria and are the progenitors of yolk cells. (**A-D**) Maximum-intensity projections of confocal sections showing dFISH of *klf4* (green) with *nanos* (**A**), or vitellaria markers *CPEB1* (**B**), *surfactant b* (**C**), and *MX1* (**D**) (magenta) in the ventral posterior region of sexually mature planarians. Dashed line denotes planarian boundary. (**A’-D’**) Single confocal sections of dFISH corresponding to **A-D**. (**A’**) dFISH of ventrally expressed *klf4* (green) and *nanos* (magenta). Almost all *klf4^+^* cells co-express *nanos* whereas *klf4* is expressed in a subset of *nanos^+^* cells. (**B’-D’**) *klf4* is expressed in a subset of *CPEB1^+^* (**B’**) and *surfactant b^+^* (**C’**) yolk cells, but not in *MX1^+^* yolk cells (**D’**). (**A’-D’**) Side panels are high-magnification views of outlined areas showing *klf4* single- (white arrowheads) and double-positive cells (yellow arrowheads). Note the increase in cell size as *klf4^+^* cells differentiate into *CPEB1^+^*, *surfactant b^+^*, and ultimately *MX1^+^* yolk cells. (**E**) Schematic depicting genes expressed during developmental progression of yolk cell lineage. (**F-G**) Maximum-intensity projections of confocal sections showing FISH of *nanos* (**F**) and *MX1* (**G**) (green) in ventral tail region of control and *klf4* RNAi animals. Dashed line denotes planarian boundary. N=3 experiments, n=8-15 planarians. *klf4* RNAi results in loss of *nanos*-expressing cells and a reduction of *MX1^+^* yolk cells in the vitellaria. (**A’-D’, F-G**) Nuclei are counterstained with DAPI (gray). Scale bars, 200 µm (**A-D, F-G**), 50 µm for overview images, 20 µm for side panels (**A’-D’**).

Taken together, these results are consistent with a model in which *klf4^+^*/*nanos^+^* cells define the origin of the yolk cell lineage and that *nanos*, *CPEB1*, *surfactant b*, and *MX1* are expressed in a partially overlapping, stepwise fashion as yolk cells differentiate (Fig 5E).

To characterize the developmental origins of the vitellaria, we examined the expression patterns of *klf4*, *nanos*, *CPEB1*, *surfactant b*, and *MX1* by dFISH at two earlier stages of planarian development. One-week-old hatchlings did not express any of these yolk cell markers in the presumptive vitellarial regions (n=0/40 hatchlings) (S4A Fig). Later in development, 100% of 2- to 3-week-old juveniles (n=26/26 planarians) possessed ventrally located *klf4^+^* cells in the region where vitellaria form (S4B Fig). Additionally, all juvenile worms expressed *nanos* (n=7/7), *CPEB1* (n=6/6) and *surfactant b* (n=7/7) in their vitellaria, but few expressed *MX1* (n=2/6) (S4B Fig). Therefore, *klf4^+^*/*nanos^+^* yolk-cell progenitors are specified post-embryonically (after hatchlings have developed into juvenile planarians), and these cells continue to differentiate into *CPEB1^+^*, *surfactant b^+^*, and ultimately *MX1^+^* yolk cells as planarians mature into adults.

Is *klf4* also required for the development of the yolk cell lineage? We induced *klf4* RNAi beginning in hatchlings (8-12 feedings) and examined the effects on early and late yolk cells by FISH to detect *nanos* and *MX1.* Knockdown of *klf4* resulted in a dramatic reduction of all yolk cells (Fig 5F and 5G), confirming the requirement for *klf4* in the development of the yolk cell lineage and suggesting that this ventral *klf4^+^*/*nanos^+^* population is indeed the progenitor of yolk cells.

### Yolk cells share features with neoblasts and germ cells

Our results suggest that *klf4* marks the top of both germ cell and yolk cell lineages. Yolk cells are technically somatic since they do not generate gametes, yet it has long been postulated that flatworm yolk cells may share an evolutionary origin with oocytes (the female germline) [44]. One hypothesis is that yolk cells were derived from germ cells in the course of evolution and that a split/divergence between these two cell types may have occurred in the common ancestor of all ectolecithal flatworms [47–49]. As we found that both yolk cells and the female germline share *klf4* and *nanos* expression, we wondered whether yolk cells share other germ cell characteristics, such as expression of *piwi-1* and *germinal histone H4* (*gH4*) (S5A Fig), two transcripts thought to be expressed exclusively in neoblasts and germ cells [34,43,50–53]. By dFISH, we found that the vast majority of *klf4^+^* cells in the vitellaria are also *piwi-1^+^* (94%, n=789 *klf4^+^* cells) and *gH4^+^* (98%, n=399 *klf4^+^* cells) (Fig 6A and 6E, S5B and S5C Fig). Unlike other somatic tissues, in which *piwi-1* mRNA is degraded during differentiation [26], *piwi-1* expression perdures during yolk cell differentiation and is still detected in most *CPEB1^+^* cells (80%, n=2801 *CPEB1^+^* cells) and *surfactant b^+^* cells (55%, n=2136 *surfactant b^+^* cells), but not in *MX1^+^* cells (0%, n=284 *MX1^+^* cells) (Fig 6B–6D). Similarly, *gH4* is co-expressed in most *surfactant b^+^* cells (64%, n=4410 *surfactant b^+^* cells) (Fig 6F and S5D Fig). Thus, similar to the germ cell lineages in testes and ovaries, *piwi-1* and *gH4* expression persist in differentiating yolk cells.

**Fig 6.**
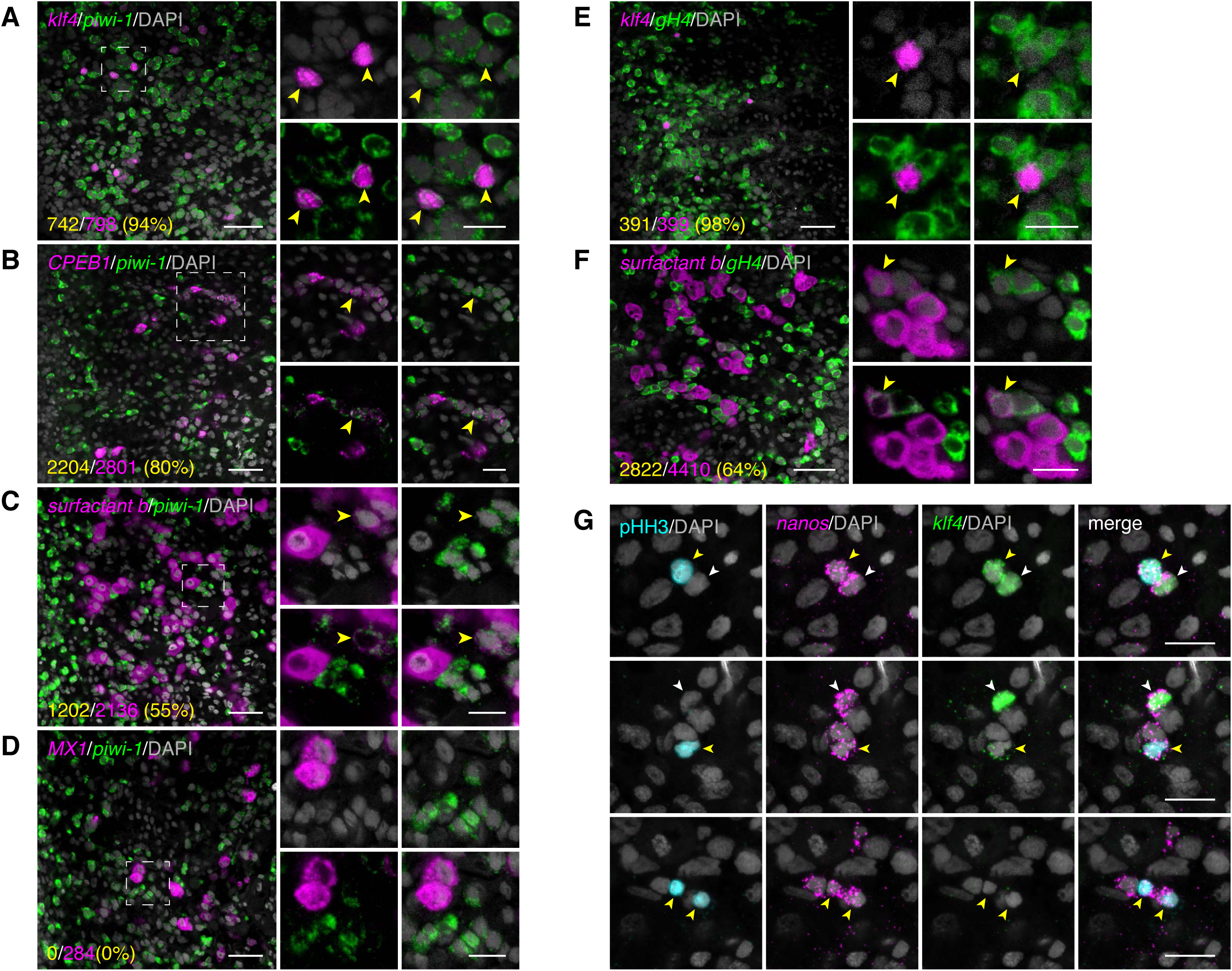
Yolk cells share features with neoblasts and germ cells. (**A-D**) Single confocal sections showing dFISH of neoblast and germ cell marker *piwi-1* (green) and *klf4* (**A**), *CPEB1* (**B**), *surfactant b* (**C**), and *MX1* (**D**) (magenta). Side panels are high-magnification views of outlined areas showing *piwi-1* double-positive cells (yellow arrowheads). (**E-F**) Single confocal sections showing dFISH of neoblast and germ cell marker *gH4* (green) and *klf4* (**E**) and *surfactant b* (**F**) (magenta). Side panels are high-magnification views of outlined areas showing *gH4* double-positive cells (yellow arrowheads). (**G**) Maximum-intensity projections of confocal sections (5 µm thick) imaged from the ventral posterior region of sexually mature planarians showing *klf4* (green) and *nanos* (magenta) dFISH with pHH3 (cyan) immunostaining in vitellaria. *klf4^+^*/*nanos^+^* vitellocytes with high (top panels) and low levels (middle panels) of *klf4* expression are mitotically active. *klf4^−^*/*nanos^+^* yolk cell progenitors are able to divide (bottom panels). (**A-G**) Nuclei are counterstained with DAPI (gray). Scale bars, 50 µm for overview images, 20 µm for side panels (**A-F**), 20 µm (**G**).

In addition to the retention of germ cell features in yolk cells, these cells are mitotically active. We detected PHH3 staining in *klf4^+^*/*nanos^+^* as well as *klf4^−^*/*nanos^+^* yolk cells (Fig 6G). Taken together, these results show that even though yolk cells do not give rise to gametes (and are therefore not germ cells), they do exhibit several germ cell characteristics, including expression of the germline markers *nanos, piwi-1,* and *gH4*, and the capacity to proliferate.

### Vitellaria contain distinct cell types: a yolk cell lineage and non-yolk support cells

Gonads are not composed solely of germ cells: they also contain somatic support cells (or niche cells) that govern germ cell behavior. Thus, we asked whether vitellaria contain non-yolk vitelline support cells and whether they could play a niche-like role in maintaining the *klf4^+^*/*nanos^+^* stem/progenitor population for sustaining the yolk cell lineage. Previously, expression of the orphan G-protein coupled receptor *ophis*, a somatic gonadal cell marker, was detected in the vitellaria, but its role there was not characterized [39]. We found that in mature sexual planarians, the vitellaria are arranged in an extensively branched network containing two populations of *ophis*-expressing cells: *ophis^high^* cells, which express *ophis* predominantly in the nucleus, and *ophis^low^* cells with weak signal throughout the cell (Fig 7A). *ophis^+^* cells are interspersed throughout the vitellaria, similar to *klf4^+^* cells (S6A-S6D Fig). *klf4^+^* cells are tightly juxtaposed with *ophis^high^* cells, however, they never co-express high levels of *ophis* (0% *klf4^+^* cells are *ophis^high^*, n=368 *klf4^+^* cells) (Fig 7A). On the other hand, a large fraction of *klf4^+^* cells are *ophis^low^* (60% *klf4^+^* cells are *ophis^low^*, n=368 *klf4^+^* cells). These results led us to hypothesize that *ophis^low^* versus *ophis^high^* cells represent two distinct classes of cells in the vitellaria: *ophis^low^* cells constitute the yolk cell lineage proper and *ophis^high^* cells are support cells that closely associate with the yolk cells and comprise the remaining structure of the vitellaria.

**Fig 7.**
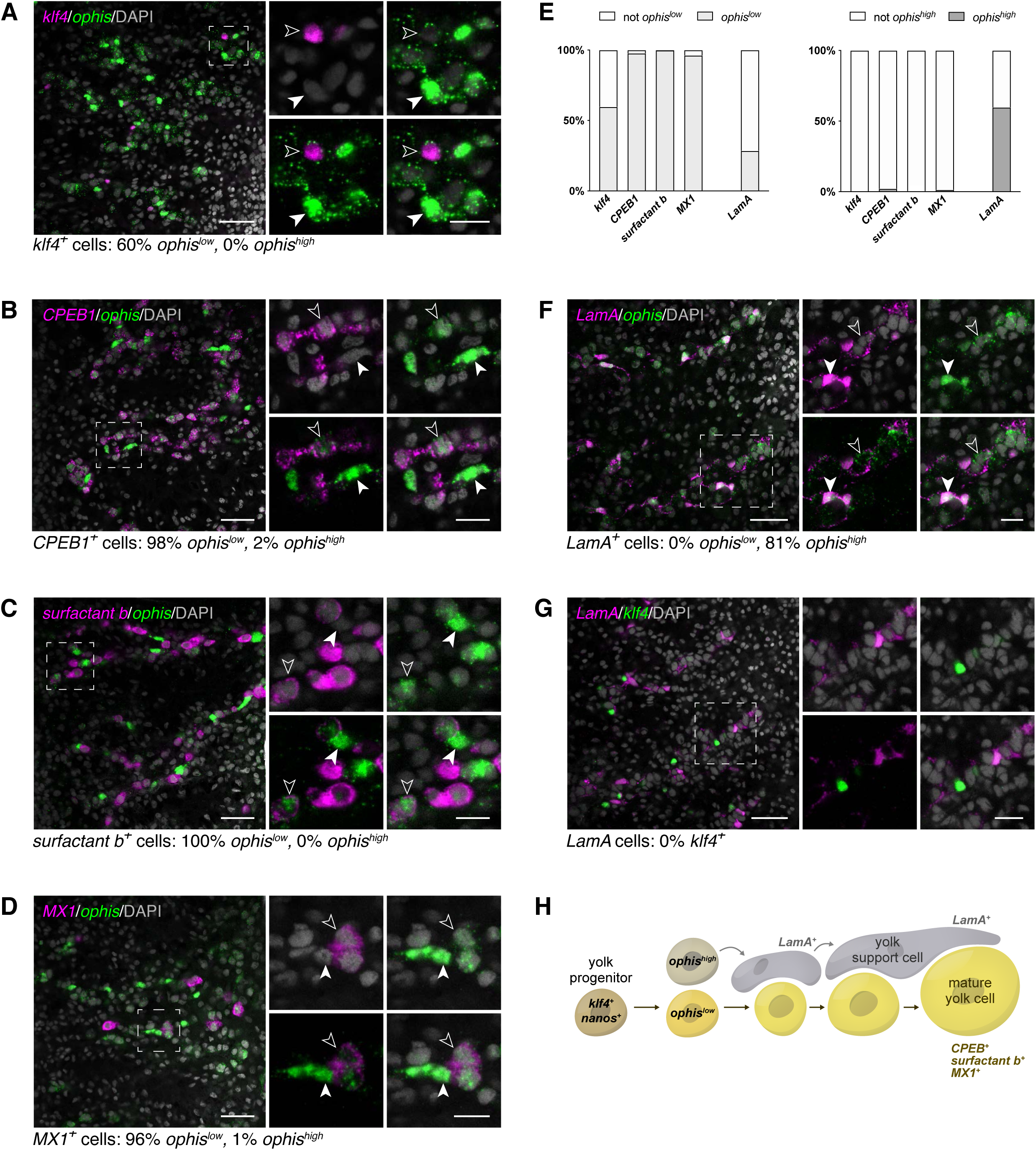
Vitellaria contain distinct cell types: yolk cells and non-yolk support cells. (**A-D, F-G**) Single confocal sections showing dFISH. Side panels are high-magnification views of outlined areas. (**A**) dFISH of *klf4* (magenta) and vitellaria marker *ophis* (green). *ophis^high^* cells do not co-express *klf4* (filled white arrowhead) but *ophis^low^* cells do (unfilled white arrowhead). (**B-D**) dFISH of *ophis* (green) and *CPEB1* (**B**), *surfactant b* (**C**), and *MX1* (**D**) (magenta). *ophis^low^* cells express yolk cell lineage differentiation markers. (**E**) proportion of cells in the vitellaria that co-express low levels (left) vs high levels (right) of *ophis*. *ophis^low^* cells predominantly co-express markers of the yolk cell lineage. Conversely, most *ophis^high^* cells co-express *LamA* but do not express yolk cell markers. (**F**) dFISH of *LamA* (magenta) and *ophis* (green). *ophis^high^* cells co-express *LamA* (filled white arrowhead) whereas *ophis^low^* cells do not (unfilled white arrowhead). (**G**) dFISH of *LamA* (magenta) and *klf4* (green). *LamA* and *klf4* are never co-expressed in the same cells. (**A-D, F-G**) Nuclei are counterstained with DAPI (gray). Scale bars, 50 µm for overview images, 20 µm for side panels. (**H**) Schematic depicting genes expressed during developmental progression of *ophis^low^* yolk cells and associated *ophis^high^* support cells.

If the *ophis^low^* population represents the yolk cell lineage of which *klf4^+^* cells are the precursors, then we would expect *klf4^−^*/*ophis^low^* cells to express markers of progressive yolk cell differentiation. Consistent with this idea, almost all *CPEB1^+^*, *surfactant b^+^*, and *MX1^+^* cells co-express low levels of nuclear and cytoplasmic *ophis* mRNA (98%, n=2914 *CPEB1^+^* cells; 100% n=2015 *surfactant b^+^* cells; 96%, n=256 *MX1^+^* cells) (Fig 7B-7E). On the other hand, high levels of nuclear *ophis* were rare in *CPEB1^+^*, *surfactant b^+^*, and *MX1^+^* cells (2%, n=2914 *CPEB1^+^* cells; 0%, n=2015 *surfactant b^+^* cells; 1%, n=256 *MX1^+^* cells). These results indicate that *ophis^low^* expression emerges in a subset of *klf4^+^* yolk cell progenitors and subsequently persists as these cells differentiate (Fig 7H), whereas *ophis^high^* expression defines a distinct cell type in the vitellaria.

In agreement with the model that *ophis^high^* cells constitute a separate cell lineage, the majority of these cells do not express yolk cell markers (0%, n=784 *ophis^high^* cells are *klf4^+^*; 12%, n=519 *ophis^high^* cells are *CPEB1^+^*; 15%, n=573 *ophis^high^* cells are *surfactant b^+^*; 1%, n=521 *ophis^high^* cells are *MX1^+^*). Instead, most *ophis^high^* cells express *Laminin A* (*LamA*) (81%, n=440 *ophis^high^* cells are *LamA^+^*) (Fig 7F), a gene expressed in the vitellaria (S6E and S6F Fig) as well as in somatic gonadal cells in the testes and ovaries (S6G Fig). This result corroborates the finding that *ophis^high^* expression marks support cells within the vitellaria. Notably, *klf4* and *LamA* are never co-expressed within the vitellaria (0%, n=540 *klf4^+^* cells, 0%, n=867 *LamA^+^* cells) (Fig 7G). Taken together, our data suggest that two cell lineages exist in the vitellaria: the yolk cell lineage (*ophis^low^*) which includes *klf4^+^* cells, and a second population made up of *ophis^high^*/*LamA^+^* cells. It was previously reported that *ophis* transcript was expressed in the somatic gonadal cells of the ovary [39]. In addition to this expression pattern, we detect low levels of *ophis* expression in the oogonial lineage, similar to yolk cells (S6H Fig). The dichotomy between *ophis^low^* versus *ophis^high^* expression in the germline and somatic lineages of the ovary is reminiscent of what we observed in the two vitellarial lineages.

### Gonadal niche factor *ophis* is required to maintain the yolk cell lineage

Previous work has shown that *ophis* is required for proper development of both male and female gonads in planarians [39]. To address whether *ophis* is a shared molecular regulator of gonads and vitellaria, we performed RNAi knockdown of *ophis* in hatchlings until they reached sexual maturity and analyzed the effects on the vitellaria by FISH (Fig 8A-8C). *ophis* knockdown resulted in a dramatic loss of the *LamA^+^* cells throughout the vitellaria, but did not affect *LamA* expression within the gut (Fig 8A). We also observed a significant reduction of *klf4^+^* cells and complete loss of mature *MX1^+^* yolk cells in *ophis* RNAi animals (Fig 8B and 8C). Although we cannot distinguish the functions of *ophis^high^* from *ophis^low^* cells in the vitellaria by available techniques in planarians, it is clear from our data that *ophis* is essential for the maintenance of support cells (*ophis^high^*/*LamA^+^*) in the vitellaria and is required (perhaps through the action of support cells) for the maintenance and differentiation of *klf4^+^* yolk cell progenitors.

**Fig 8.**
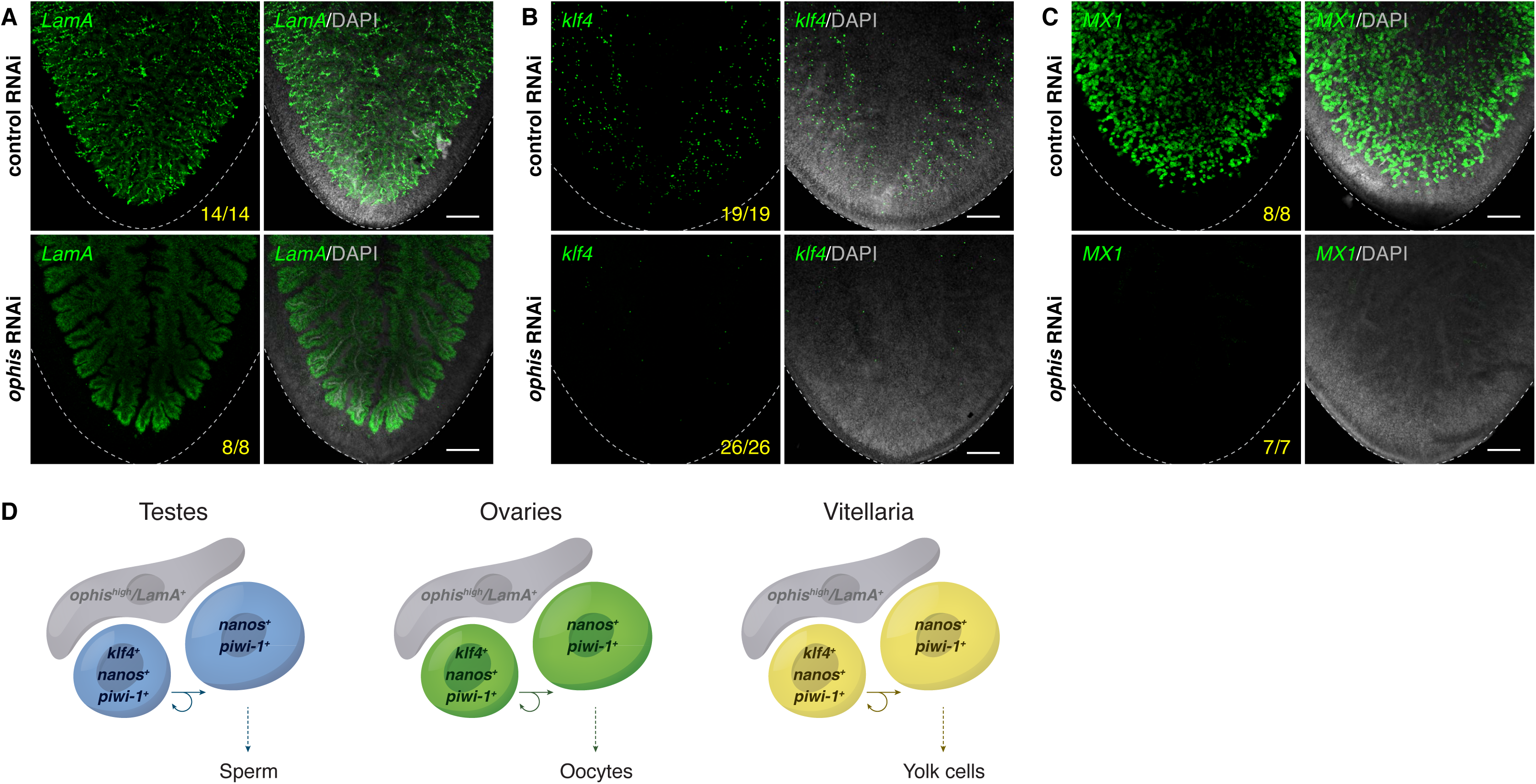
Germ cell niche factor *ophis* is required to sustain yolk cell production/ vitellogenesis. (**A-C**) Maximum-intensity projections of confocal sections showing FISH of *LamA* (**A**), *klf4* (**B**), and *MX1* (**C**) (green) in the ventral posterior region of sexually mature control vs *ophis* RNAi animals. Dashed line denotes planarian boundary. N=3-5 experiments, n=7-26 planarians. (**A**) *ophis* RNAi results in a dramatic loss of the *LamA^+^* cells throughout the vitellaria. Note that *LamA* expression is only visible in the branched gut in *ophis* RNAi planarians. (**B-C**) *ophis* RNAi results in a reduction of *klf4^+^* yolk cell progenitors and *MX1^+^* differentiated yolk cells. (**A-C**) Nuclei are counterstained with DAPI (gray). Scale bars, 200 µm. (**D**) Model depicting similarities shared between gonads (where gametogenesis occurs) and vitellaria (where yolk cell production occurs). *klf4^+^*/*nanos^+^*/*piwi-1^+^* GSCs in testes and ovaries divide and give rise to *klf4^−^* /*nanos^+^*/*piwi-1^+^* progeny. These germ cells are supported by *ophis^+^* somatic gonadal niche cells. Vitellaria are comprised of *klf4^+^*/*nanos^+^*/*piwi-1^+^* “germ-cell-like” yolk progenitors that are mitotically competent, sustain yolk cell production, and are supported by *ophis^high^* support cells.

## Discussion

Most animals specify PGCs and segregate them from somatic tissues only once, early in development. Within developed gonads, germ cells are generated from GSCs for the reproductive life of the organism. Planarians also specify PGCs in development but are able to continuously regenerate new germ cells from pluripotent stem cells throughout their lifetime. Whether or not planarians also maintain GSCs is less clear, especially since theoretically they could reseed gonads with new germ cells from their somatic stem cells (neoblasts) throughout adulthood. Characterizing the regulators that define planarian germ cells and function in their specification and maintenance will reveal important clues for understanding the remarkable ability of planarians to faithfully regenerate germ cells.

### Elucidating early stages of the germ cell lineage

We found that *klf4* expression marks the earliest/least differentiated germ cell state in planarians. Early hatchlings specify PGCs that co-express *klf4* and *nanos* dorsolaterally, where adult testes will ultimately reside. Thus, male PGCs likely do not undergo extensive migration to the somatic gonad. Instead, they are likely to be specified along with *dmd1*-expressing somatic gonadal niche cells, and then differentiate in situ as the testis grows/elaborates during reproductive maturation. As hatchlings develop into juveniles, testis primordia grow in size and two successive populations of *klf4^−^* germ cells arise (*nanos^+^* and *nanos^−^*). These populations emerge as testes develop, strongly suggesting that they represent the first step in germ cell differentiation and that cessation of *klf4* expression may be required for germ cell differentiation to proceed.

We observed that *klf4* is expressed in virtually all *nanos^+^* cells in early hatchlings, and then becomes increasingly restricted to a smaller subset of *nanos^+^* cells as planarians sexually mature. The restriction of *klf4* expression to a subset of *nanos^+^* germ cells holds true in asexual planarians as well, where the number of germ cells in both gonadal primordia increases as animals grow (Sato et al., 2006). Small asexual planarians express *klf4* in almost all *nanos^+^* germ cells, whereas larger asexuals have proportionally fewer double-positive cells in their testis primordia. Our results refine the stage at which development arrests in asexuals: in growing asexuals, *klf4^+^*/*nanos^+^* cells can only carry out the first step of development (into *klf4^−^*/*nanos^+^* cells), further reinforcing the idea that the germ cell lineage progresses in this direction.

Additionally, we have shown that *klf4* is required for the specification of germ cells. RNAi knockdown of *klf4* in soma-only head fragments results in regenerated animals that do not re-specify *nanos^+^* germ cells, even though new testis somatic gonadal support cells (*dmd1^+^*) that form the niche are made. Our data indicate that *klf4* is required cell autonomously for the de novo specification of germ cells. Taken together, these observations support a model in which *klf4* expression marks the top of the germ cell hierarchy and that expression of *klf4* is required for the acquisition of germ cell fate.

Although animals specify their germline in different ways (preformation vs induction), a conserved feature of newly specified PGCs is the repression of somatic differentiation transcriptional programs. Posttranscriptional regulation through the action of conserved germline-specific RNA regulators such as *vasa*, *pumilio*, *nanos*, and *piwi* plays an outsized role in controlling germ cell fate, survival, proliferation, and differentiation. Germ cell fate specification at the transcriptional level is less well understood. During mouse embryogenesis, PGCs are specified from pluripotent epiblast cells by BMP signals from the extraembryonic ectoderm and the visceral endoderm through the action of Smads [54–57]. Critical regulators of PGC specification have been described, including transcription factor genes *Prdm1* (which encodes BLIMP1) and *Prdm14* [58–65]. A key role of BLIMP1 is to induce expression of *Tcfap2c* (which encodes the transcription factor AP2γ) [64,66,67], and together, BLIMP1, PRDM14, and AP2γ are important for initiating PGC specification, repressing expression of somatic genes, activating expression of PGC-specific genes, and driving epigenetic reprograming [62,66–69].

Recent studies on emerging models have shown that some of the molecular mechanisms regulating PGC specification may be conserved. In the cricket *Gryllus bimaculatus*, PGCs are specified in response to BMP signaling via the action of Blimp-1 [70, 71]. Additionally, in the cnidarian *Hydractinia symbiolongicarpus*, a homolog of AP2 is an inducer of germ cell fate [11]. Although the inductive cues that control germ cell fate in *S. mediterranea* remain to be identified, here we identify a transcription factor, Klf4, required for germ cell specification. It is worth noting that Klf4 is a crucial pluripotency factor in mammals. Furthermore, pluripotency genes *Oct4*, *Sox2*, and *Nanog* are expressed in mouse PGCs [72], reflecting the importance of maintaining pluripotency in germ cells. Therefore, future identification of Klf4 targets in *S. mediterranea* will not only elucidate the transcriptional program required for promoting germ cell fate from pluripotent neoblasts but may also provide important clues into how germline pluripotency is maintained.

### Are *klf4*-expressing cells true stem cells?

GSCs are characterized by the ability to undergo self-renewing divisions in which one daughter remains a stem cell and the other differentiates. Consistent with the hypothesis that *klf4^+^* cells are GSCs, *klf4*-expressing cells in both testes and ovaries are mitotically active throughout post-embryonic development. However, technical limitations precluded us from testing whether mitotic *klf4^+^* cells undergo self-renewing divisions. Alternatively, it is possible that no resident GSC population exists within the gonads themselves; instead neoblasts residing in the gonad-adjacent parenchyma may be continually specified as new germ cells that then differentiate directly without self-renewing. Either way, dFISH experiments with *klf4* and *nanos* have uncovered heterogeneity within the early male and female germ cell compartments. Furthermore, developmental timeline experiments have allowed us to define the early germ cell lineage with *klf4^+^*/*nanos^+^* germ cells at the top of the hierarchy. Prolonged inhibition of *klf4* via RNAi during post-embryonic development and sexual maturation led to a dramatic loss of early germ cells in both testes and ovaries, resulting in agametic gonads in some animals. This result suggests that *klf4* is required for the maintenance of GSCs (or germ cell lineal progenitors) to sustain gametogenesis. Future experiments will explore whether *klf4^+^*/*nanos^+^* cells represent true GSCs and whether this newly defined lineage progresses in a unidirectional manner, or if all *nanos*-expressing cells retain GSC-like potential.

### Bidirectional soma-germ cell communication in the ovary

Intriguingly, loss of germ cells in the ovary led to a corresponding increase in ovarian somatic gonadal cells (*ophis^+^* and *LamA^+^*). This result reveals that soma-germline communication in the planarian ovary is bidirectional. The importance of somatic support cells for germ cell development is undisputed. However, far less is known about how germ cells signal back to their somatic microenvironment [73–75]. In planarians, somatic cell expansion in the ovary in response to germ cell loss suggests that somatic and germ cell numbers are coordinated via a feedback mechanism. What signals regulate this feedback loop? How does the planarian ovary balance somatic and germ cell numbers to achieve an equilibrium between these two cell types? The planarian ovary presents a unique opportunity to investigate the mechanisms involved in soma-germline coordination during development, homeostasis, and regeneration.

While both gonads contain *klf4^+^*/*nanos^+^* putative GSCs, there is also a population of these cells anterior to each ovary. They may be germ cell progenitors that migrate posteriorly and enter the ovary, where they then give rise to *nanos^−^* oogonia/oocytes. Alternatively, *klf4^+^*/*nanos^+^* cells may be specified in a permissive zone along the medial-posterior regions of the cephalic ganglia, but only the posterior-most germ cells located at the base of the brain (where the somatic gonad is located) are then able to associate with somatic gonadal cells and consequently instructed to differentiate. Until we are able to specifically ablate this population, its contribution to the ovary (or lack thereof) will remain mysterious.

### A shared evolutionary origin of germ cells and yolk cells?

A unique reproductive feature of flatworms is ectolecithality: a developmental novelty in which oocytes develop with little/no yolk while specialized yolk cells are produced ectopically. For embryogenesis to occur, the fertilized oocyte and numerous yolk cells must be deposited together in egg capsules. As yolk cells are the sole source of embryonic nutrients, ectolecithality has led to marked evolutionary and functional consequences on embryonic development. For example, yolkless embryos develop temporary organs (e.g., embryonic pharynx, primitive gut) that facilitate uptake of maternally provided yolk/nutrients early in embryogenesis [76].

Recent phylogenetic analyses have shed light on the origin of ectolecithality in flatworms. One group of flatworms produces oocytes and yolk cells within a single organ (the germovitellarium); another group partitions egg- and yolk cell-production into two distinct organs (the germarium/ovary and vitellaria). This latter group is known as Euneoophora and includes planarians and parasitic flatworms. Although traditional phylogenies grouped both types of ectolecithal worms together, recent phylogenies suggest that they evolved independently [47–49]. Thus, the ectolecithal common ancestor of all euneoophorans likely evolved from more primitive endolecithal (“yolky egg”-producing) flatworms [48, 49], consistent with a model in which yolk cells in ectolecithal flatworms evolved from ancestral “yolky” germ cells. These phylogenetic studies recognized that molecular similarities between germ cell and yolk cell precursors would lend further support to the shared evolutionary origin hypothesis [47–49]. Here we provide molecular and developmental evidence suggesting that yolk cells and germ cells are homologous. Even though yolk cells do not produce gametes and, therefore, are not de facto germ cells, they share several molecular and cellular characteristics in common with germ cells (Fig 8D). Yolk cells express both *klf4* and *nanos*: two markers that define male and female germ cell lineages. Similarly to testes and ovaries, *klf4* expression in vitellaria is restricted to a subset of *nanos^+^* yolk cells, suggesting that *klf4^+^*/*nanos^+^* cells define the lineal progenitors of yolk cells. We also find that yolk cells express *piwi-1* and *gH4*, which until this work, were reported to be expressed exclusively in neoblasts and germ cells. *piwi-1* and *gH4* are highly expressed in neoblasts but downregulated in their immediate somatic progeny. In contrast to the soma, but similar to *piwi-1* and *gH4* expression in male and female germ cell lineages, expression of these genes is sustained in differentiating (*klf4^−^*/*nanos^−^*) yolk cells. This sustained expression of neoblast/germ cell markers provides another molecular similarity between germ cells and yolk cells.

Surprisingly, we observed mitosis in yolk cells. Previously, the only planarian somatic cells thought to have mitotic activity were neoblasts. Although yolk cells are technically somatic, our results clearly indicate that like germ cells, a subset of yolk cells is mitotically competent. The observation that both *klf4^+^*/*nanos^+^* and *klf4^−^*/*nanos^+^* yolk cells divide indicates that mitotic ability is not limited to the earliest progenitor in the yolk cell lineage. Are pHH3^+^/*klf4^+^*/*nanos^+^* cells undergoing self-renewing divisions? Do *piwi-1^+^*/*klf4^+^*/*nanos^+^* cells represent a new stem cell population in planarians? With the sole exception of planarian gonads, no other planarian organ contains a resident stem cell (or dividing cell) population. Instead, dividing neoblasts in the parenchyma are the only source of new differentiated somatic cells, which then integrate into existing tissues. The planarian vitellarium provides an intriguing case study to understand the regulation of stem cell populations in planarians.

These similarities between the female germ cell and yolk cell lineages prompted us to ask whether ovaries and vitellaria also share structural features. For example, is there a distinct lineage of somatic support cells that act as a niche? Gonads are typified by the presence of somatic support cells that associate intimately with germ cells and play crucial roles in their development. We discovered that in addition to the yolk cells, vitellaria contain a second population of cells (*ophis^high^*/*LamA^+^*) with long fingerlike projections that contact all stages of the yolk cell lineage. Both *ophis* and *LamA* are also expressed in the somatic gonadal cells of the ovary. *ophis* RNAi leads to loss of *LamA^+^* vitellaria cells, a dramatic decrease in *klf4^+^* yolk cell progenitors, and a complete failure of vitellogenesis, suggesting that the *ophis^high^*/*LamA^+^* cells could function as a niche required to maintain the yolk cell stem/progenitor population. Because a significant number of *klf4^+^* yolk cell progenitors co-express low levels of *ophis*, we cannot yet distinguish definitively between a cell-autonomous versus non-autonomous role for *ophis* in yolk cell development. However, since *ophis* RNAi results in a dramatic loss of *klf4^+^* cells that far outnumbers the fraction of *klf4^+^* cells that co-express *ophis* (60%), we favor the model that *ophis* acts non-autonomously in the maintenance of *klf4^+^*/*ophis^−^* yolk cells.

Comparative analyses of gametogenesis and vitellogenesis in *S. mediterranea* have allowed us to investigate the biological phenomenon of ectolecithality and to better understand its origin in Platyhelminthes. Interestingly, *nanos* expression has been detected in early yolk cells of the parasitic flatworm *Schistosoma mansoni* [32]. Since *all* parasitic platyhelminthes (trematodes, cestodes, and monogeneans) are characterized by the presence of ectolecithality, and depend on sexual reproduction to successfully propagate, the vitellaria may provide an effective anti-helminthic target. Thus, the experimental accessibility of planarians provides an opportunity to dissect the mechanisms regulating vitellaria development, with the potential to help in the fight against their parasitic cousins.

### Conclusion

This study demonstrates the functional requirement for *klf4* in germ cell specification and maintenance in planarians, and provides evidence that *klf4* expression marks the top of the germ cell lineage. Additionally, our results suggest that *klf4* is a pivotal intrinsic regulator not only of germ cells, but also of yolk cells in a somatic reproductive structure, the vitellaria. Furthermore, we identify a new population of mitotically competent yolk cell progenitors and characterize their niche. Together, these results show that planarian germ cells and somatic yolk cells exhibit a remarkable degree of similarity, supporting the hypothesis that these two lineages share an evolutionary origin.

## Materials and Methods

### Planarian culture

Sexual *S. mediterranea* [52] were maintained in 0.75X Montjuïc salts [77] at 16-18°C. Asexual *S. mediterranea* (clonal strain CIW4) [78] were maintained in 1X Montjuïc salts at 20-22°C. Planarians were starved for one week before experimentation.

### Cloning

Target genes were cloned by PCR amplification of cDNA generated from RNA extracted from adult sexual *S. mediterranea*. Gene-specific PCR amplicons were ligated into the pJC53.2 vector via TA-cloning as previously described [51]. Anti-sense riboprobes were generated by in vitro transcription reactions with T3 or SP6 RNA polymerases [46]. dsRNA was generated using T7 RNA polymerase [79]. Sequences used for probes and dsRNA are found in S1 Table.

### In situ hybridization

FISH protocols were performed as previously described [45, 46] with the following modifications. Asexual and sexual hatchling/sexual adult planarians were killed in 7.5 % N-acetyl-L-cysteine in 1X PBS for 5/10 minutes; fixed in 4% formaldehyde in PBSTx (1X PBS + 0.1% Triton X-100) for 15/30 minutes; bleached in Bleaching Solution (1X SSC solution containing 5% deionized formamide and 1.2% hydrogen peroxide) for 2/4 hours; incubated in PBSTx containing 10 ug/ml Proteinase K and 0.1% SDS for 10/20 minutes; and re-fixed in 4% formaldehyde in PBSTx for 10/15 minutes. Planarians were blocked in Blocking Solution (5% heat inactivated horse serum, 5% Roche Western Blocking Buffer in TNTx [0.1 M Tris pH 7.5, 0.15 M NaCl, 0.3% Triton X-100]) for 2 hours at room temperature, and incubated in Blocking Solution containing anti-Digoxigenin-POD (1:2000; Roche #11207733910), anti-Fluorescein-POD (1:2000; Roche #11426346910), or anti- Dinitrophenyl-HRP (1:2000; Vector Laboratories #MB-0603) for 8 hours at 12°C. For fluorescent development of riboprobes, tyramide signal amplification (TSA) reactions were performed for 30 minutes.

### Phospho-Histone H3 immunofluorescence

Immunostaining was performed after FISH development by re-blocking planarians in Blocking Solution (5% heat inactivated horse serum, 5% Roche Western Blocking Buffer in TNTx) for 2 hours at room temperature, labeling mitotic cells with anti-phospho-Histone H3 (Ser10) (1:2000; Millipore Cat# 05-806) in Blocking Solution overnight at 12°C, washing 6X in PBSTx (30 minutes each), re-blocking for 2 hours at room temperature, and incubating with peroxidase-conjugated goat anti-mouse (1:500; Jackson ImmunoResearch Labs #115-035-044) in blocking solution overnight at 12°C. Planarians were washed 6X in PBSTx (30 minutes each) and TSA was performed for 30 minutes.

### Imaging

Confocal imaging was performed using a ZEISS LSM 880 with the following objectives: EC Plan-Neofluar 10x/0.3 M27, Plan-Apochromat 20x/0.8 M27, Plan-Apochromat 40x/1.3 Oil DIC M27. Image processing was performed using ZEISS ZEN 3.1 (blue edition) for linear adjustments and maximum intensity projections.

### RNA interference

In vitro dsRNA synthesis was performed as previously described [79] by in vitro transcription from PCR amplicons flanked by T7 promoters. In vitro transcription reactions were carried out overnight at 31°C, DNase-treated, brought up to 80-100 µl final volume with water, and annealed. dsRNA was added to liver (1:2–1:5) and fed to animals. dsRNA generated from the *CamR* and *ccdB*-containing insert of the pJC53.2 vector was used for all control RNAi feedings [51].

### Quantification and statistical analysis

Cell counting was performed manually. Counts for all experiments are detailed in S1 Data. Statistical analyses were performed using GraphPad Prism software. Statistical tests, significance levels, number of data points, planarian numbers (n), and experimental replicates (N) are provided in the text and/or figure legends.

## Acknowledgements

We thank Newmark lab members, past and present, for discussion and feedback. We especially thank Katherine Browder for excellent technical support, John Brubacher for invaluable comments on the manuscript, Tracy Chong for assistance with early characterization of *klf4*, and Matthew Stefely for providing illustrations/graphics. This work was supported by NIH R01 HD043403. M.I. was a Damon Runyon Fellow supported by the Damon Runyon Cancer Research Foundation (DRG-2135-12). P.A.N. and P.W.R. are Investigators of the Howard Hughes Medical Institute.

## Supplemental Figure Legends

**S1 Fig.**
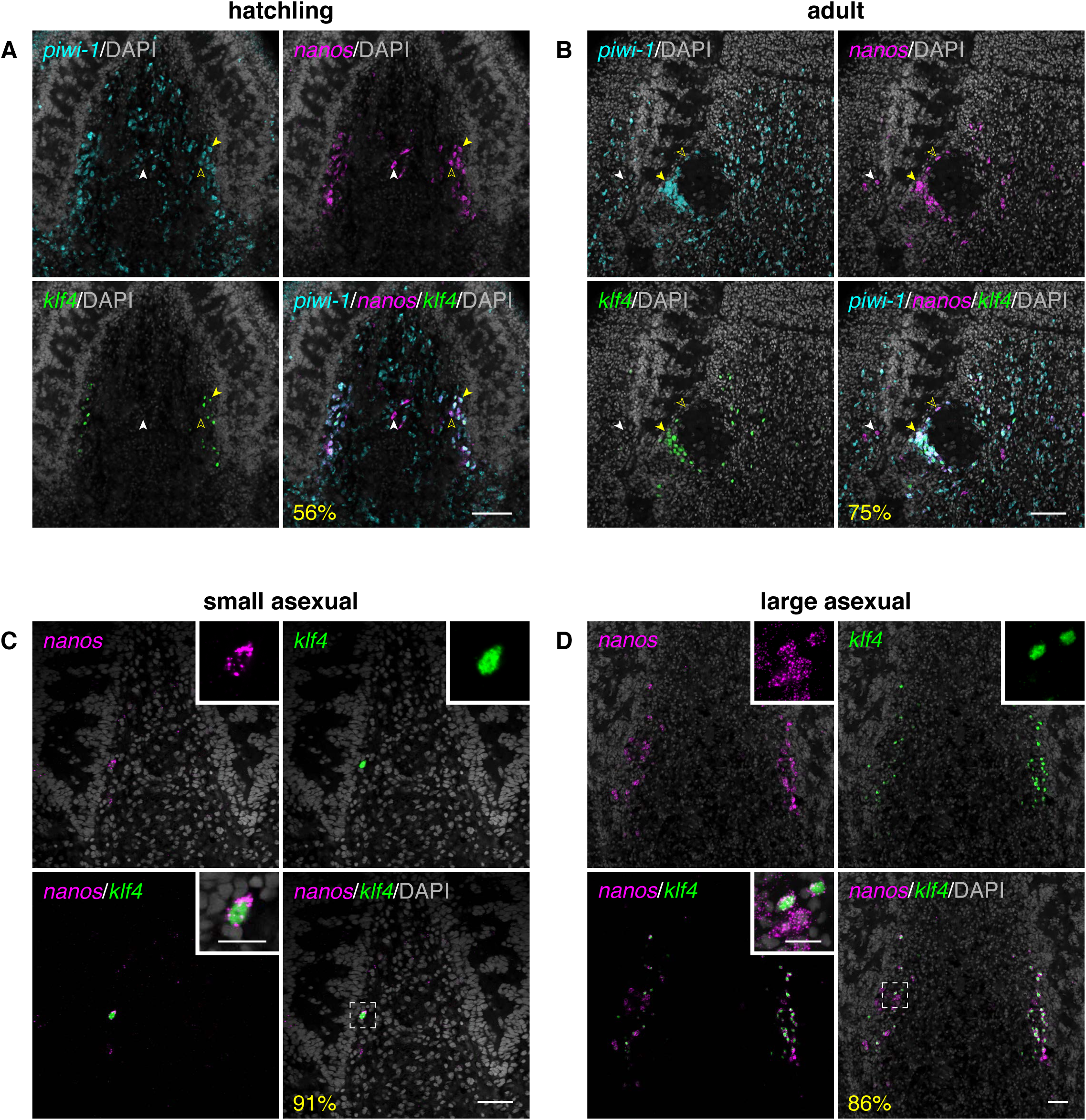
*klf4* is expressed in a subset of *nanos^+^* female germ cells in sexual and asexual planarians. (**A-B**) Confocal section showing triple FISH of *piwi-1* (cyan), *klf4* (green), and *nanos* (magenta) in female germ cells in hatchlings and sexually mature ovary. *klf4* is expressed in a subset of *nanos^+^*/*piwi-1^+^* female germ cells (compare filled (*klf4^+^*) to unfilled (*klf4^−^*) yellow arrowhead). All *klf4^+^*/*nanos^+^* germ cells are *piwi-1^+^*. A small fraction of *klf4^−^/nanos^+^* cells do not express *piwi-1* and are not germ cells (white arrowhead). (**C-D**) Confocal sections showing dFISH of *klf4* (green) and *nanos* (magenta) in female germ cells (located mediolaterally along the planarian brain) in small (**C**) and large (**D**) asexual planarians. *klf4* is expressed in a subset of *nanos^+^* female germ cells. Insets show high-magnification views of heterogeneity of *klf4* expression in *nanos^+^* cells. (**A-D**) Percentages reflect *nanos^+^* germ cells that are also *klf4^+^*. Nuclei are counterstained with DAPI (gray). Scale bars, 100 µm (**A-B**), 50 µm for whole-brain images, 10 µm for insets (**C-D)**.

**S2 Fig.**
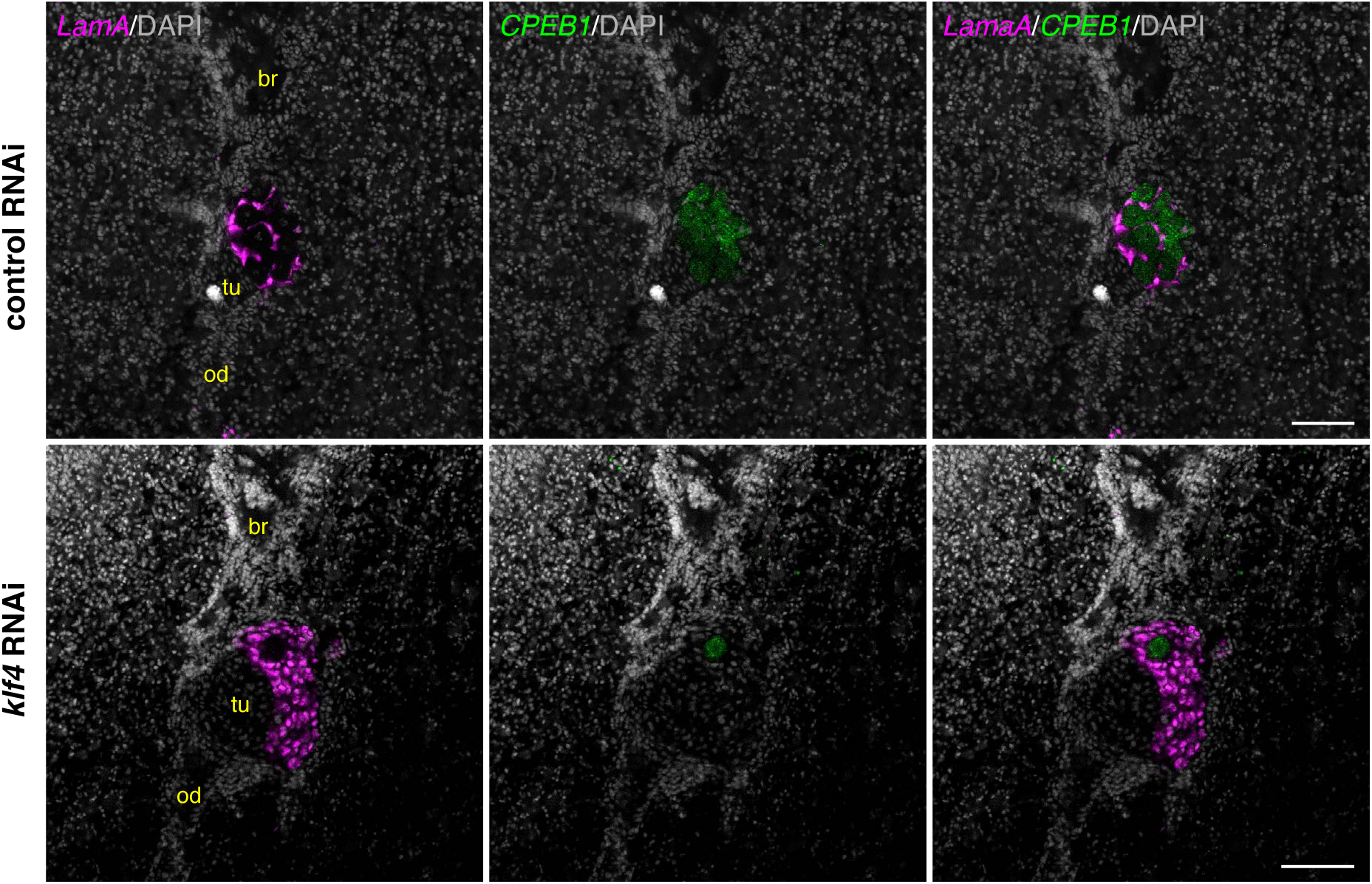
*klf4* is required for oogenesis and restricts expansion of somatic gonadal cells in adult ovaries. Single confocal section of an ovary located at the posterior of the brain (br) and anterior to the tuba/oviduct (tu/od) showing dFISH of *CPEB1* (magenta; oocytes) and *LamA* (green; somatic gonadal cells) in control and *klf4* RNAi planarians. *klf4* RNAi leads to oocyte loss and a non-autonomous increase in somatic support cells. Nuclei are counterstained with DAPI (gray). Scale bars, 100 µm.

**S3 Fig.**
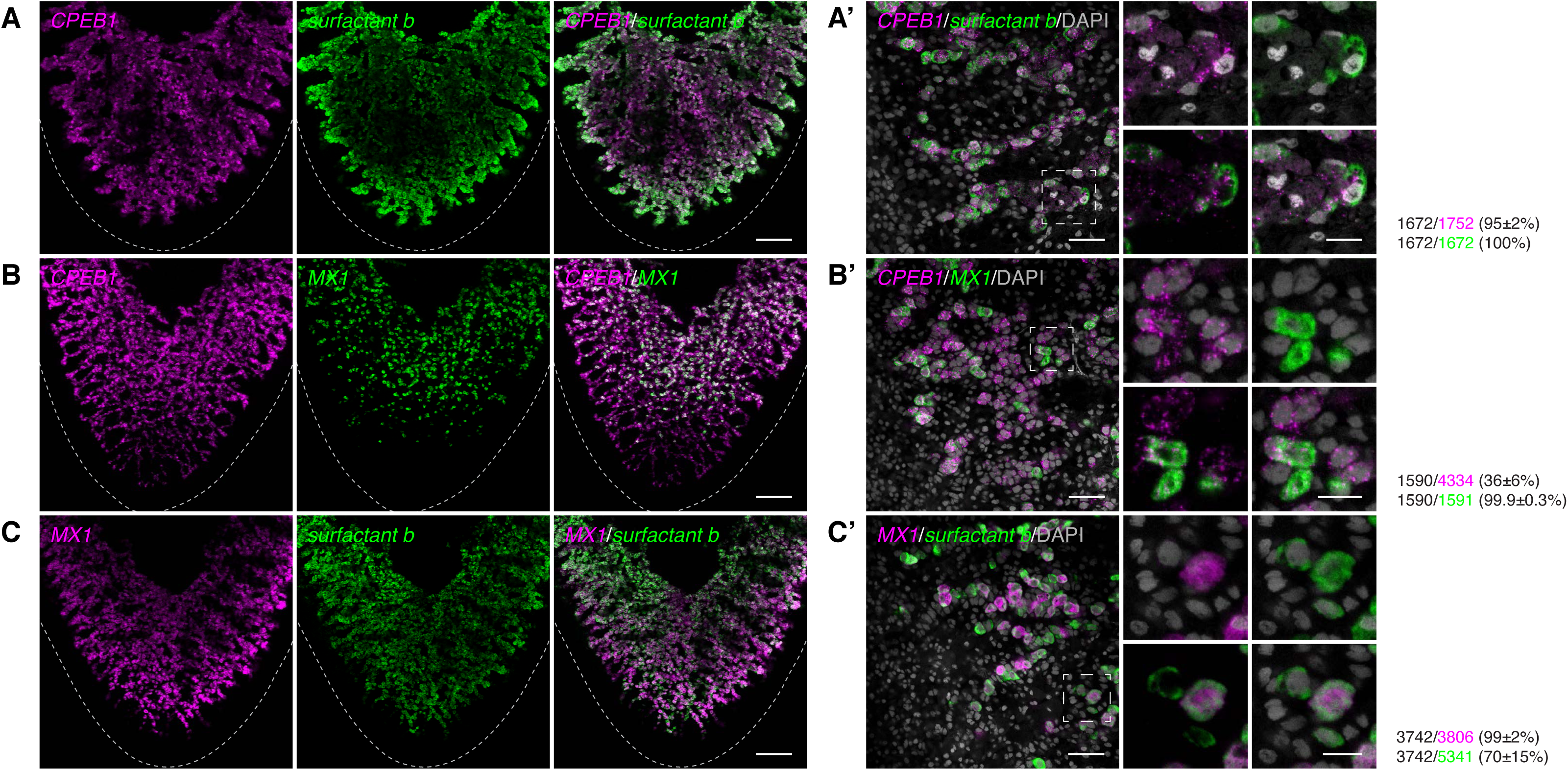
Defining the stages of yolk cell development. (**A-C**) Maximum-intensity projections of confocal sections showing dFISH of vitellaria markers *CPEB1* (**A-B**), *surfactant b* (**A, C**), and *MX1* (**B-C**) in the ventral posterior region of sexually mature planarians. Dashed line denotes planarian boundary. (**A’-C’**) Single confocal sections of dFISH corresponding to **A-C**. (**A’**) dFISH of ventrally expressed *CPEB1* (magenta) and *surfactant b* (green). Almost all *CPEB1^+^* cells co-express *surfactant b* and all *surfactant b^+^* cells are *CPEB1^+^*. (**B’**) dFISH of *CPEB1* (magenta) and *MX1* (green). A subset of *CPEB1^+^* cells co-express *MX1* whereas all *MX1^+^* cells are *CPEB1^+^*. (**C’**) dFISH of *MX1* (magenta) and *surfactant b* (green). A subset of *surfactant b^+^* cells co-expresses *MX1* whereas virtually all *MX1^+^* cells are *surfactant b^+^*. (**A’-C’**) Side panels are high-magnification views of outlined areas. (**A’-C’**) Nuclei are counterstained with DAPI (gray). Scale bars, 200 µm (**A-C**), 50 µm for overview images, 20 µm for side panels (**A’-C’**).

**S4 Fig.**
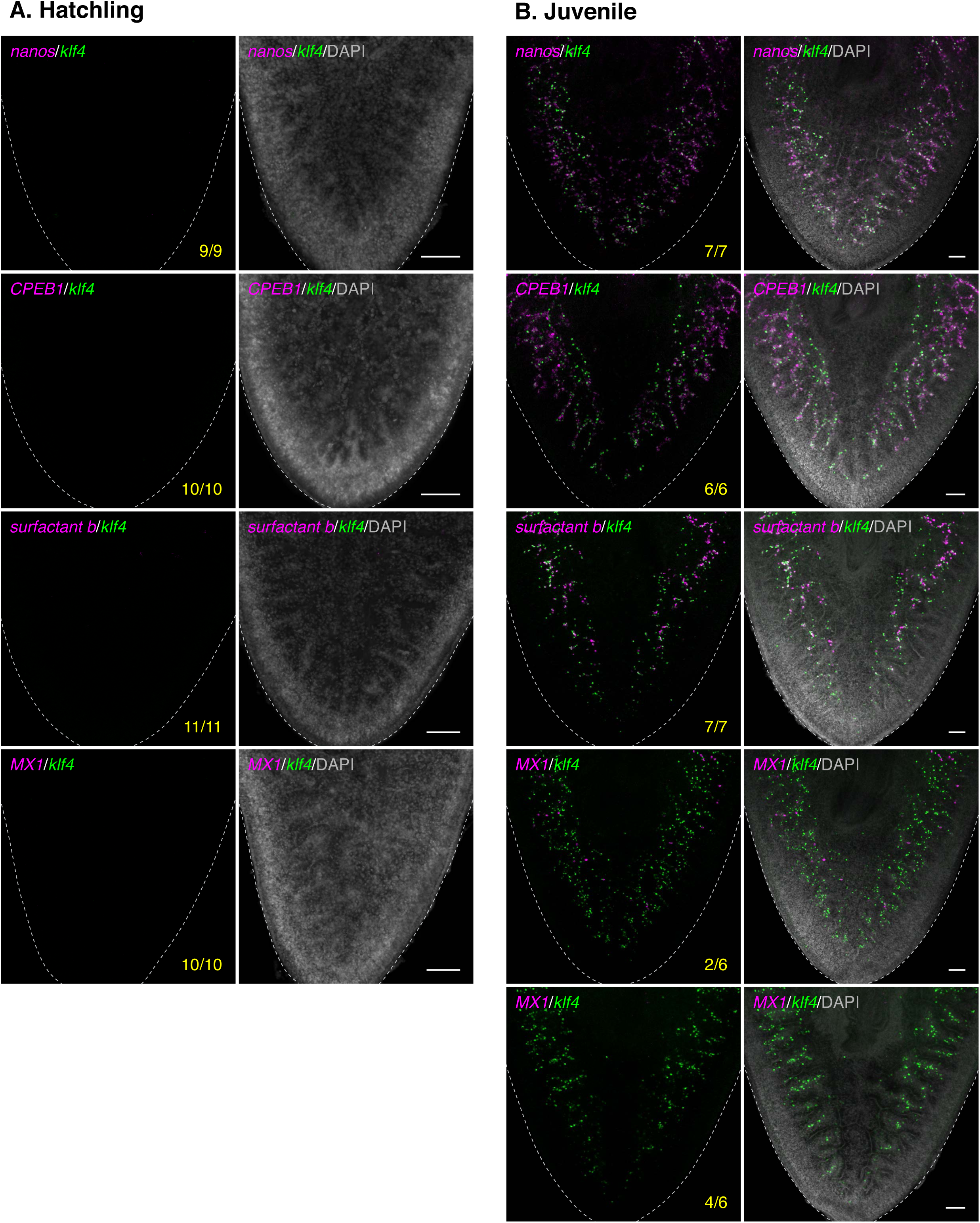
Vitellaria develop post-embryonically and produce differentiating yolk cells during sexual maturation. (**A-B**) Maximum-intensity projections of confocal sections showing dFISH of *klf4* (green) with *nanos*, or vitellaria markers *CPEB1*, *surfactant b*, or *MX1* (magenta) in the ventral posterior region of hatchlings (**A**) or juveniles (**B**). Dashed line denotes planarian boundary. (**A**) Hatchlings do not express any of the vitellaria markers tested and are devoid of vitellaria. (**B**) *klf4^+^*/*nanos^+^* yolk-cell progenitors, as well as *klf4^−^*/*nanos^+^*, *CPEB1^+^*, and *surfactant b^+^* differentiating yolk cells are detected in all juveniles. Only a fraction of juveniles express *MX1^+^* yolk cells (**B**). (**A-B**) Nuclei are counterstained with DAPI (gray). Scale bars, 100 µm.

**S5 Fig.**
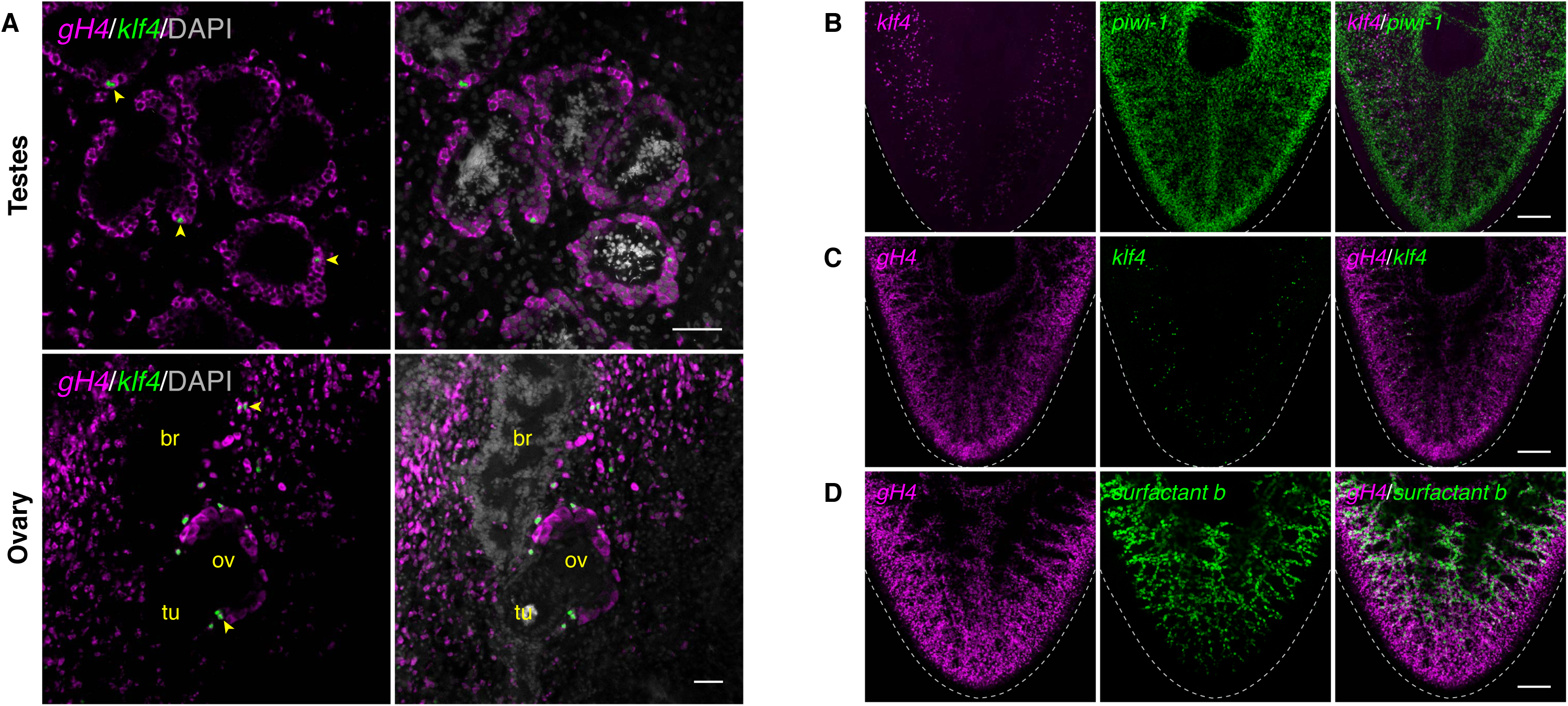
Yolk cells express neoblast/germ cell markers. (**A**) Single confocal sections showing dFISH of neoblast and germ cell marker *gH4* (magenta) and *klf4* (green). *gH4* is expressed at high levels in neoblasts as well as in spermatogonia and oogonia. *klf4^+^* cells in the testes (top panels), ovarian field, and ovary (ov) (bottom panel) co-express *gH4* (yellow arrowheads). Note the absence of *gH4* in differentiated somatic cells found in the brain (br) and tuba (tu). Nuclei are counterstained with DAPI (gray). (**B-C**) Maximum-intensity projections of confocal sections showing dFISH of *klf4* and neoblast/germline markers *piwi-1* or *gH4* in the vitellaria. (**D)** Maximum-intensity projection of confocal sections showing dFISH of *gH4* (magenta) and *surfactant b* (green). Dashed line denotes planarian boundary. Scale bars, 50 µm (**A**), 200 µm (**B-D**).

**S6 Fig.**
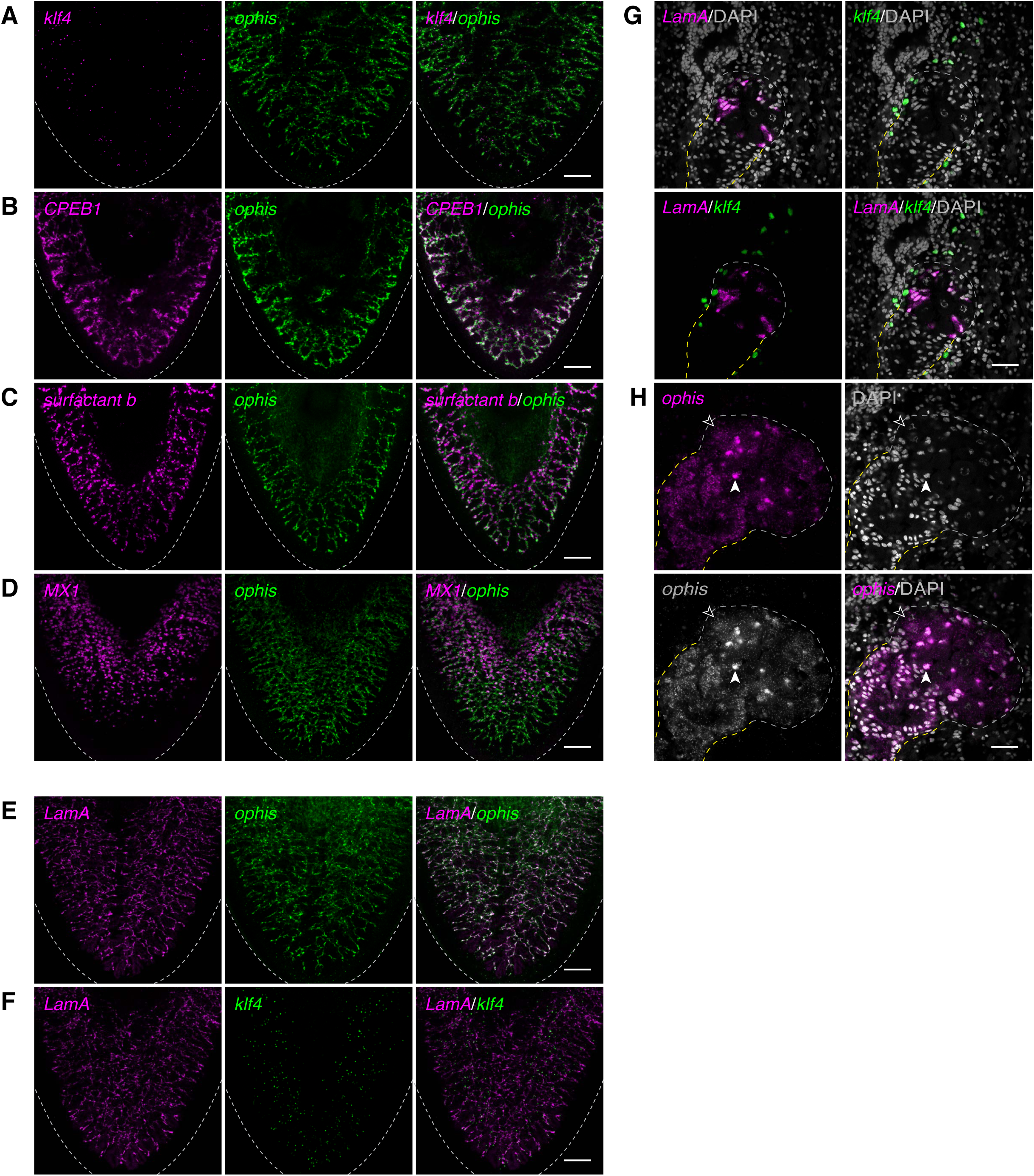
The vitellaria and ovary are comprised of two populations of *ophis*-expressing cells: *ophis^high^* versus *ophis^low^* cells. (**A-F**) Maximum-intensity projections of confocal sections showing dFISH of vitellaria markers in the ventral posterior region of sexually mature planarians. Dashed line denotes planarian boundary. (**G**) Confocal section of an ovary depicting *LamA* expression (magenta) in somatic gonadal cells and *klf4* expression (green) in early germ cells. (**H**) Confocal section of an ovary depicting *ophis^high^* expression (magenta/gray) in somatic gonadal cell nuclei (filled arrowhead) and *ophis^low^* expression in oogonia and oocytes (unfilled arrowhead). (**G-H**) Dashed line denotes ovary (white) and tuba (yellow) boundary. Nuclei are counterstained with DAPI (gray). Scale bars, 200 µm (**A-F**), 50 µm (**G-H**).

## References

1. Extavour CGM. Evolution of the bilaterian germ line: lineage origin and modulation of specification mechanisms. Integr Comp Biol. 2007;47: 770–785. doi:10.1093/icb/icm027.

2. Extavour CG, Akam M. Mechanisms of germ cell specification across the metazoans: epigenesis and preformation. Development. 2003;130: 5869–5884. doi:10.1242/dev.00804.

3. Seydoux G, Braun RE. Pathway to totipotency: lessons from germ cells. Cell. 2006;127: 891– 904. doi:10.1016/j.cell.2006.11.016.

4. Strome S, Updike D. Specifying and protecting germ cell fate. Nat Rev Mol Cell Biol. 2015;16: 406–416. doi:10.4161/worm.28641.

5. Mochizuki K, Sano H, Kobayashi S, Nishimiya-Fujisawa C, Fujisawa T. Expression and evolutionary conservation of *nanos*-related genes in *Hydra*. Dev Genes Evol. 2000;210: 591– 602. doi:10.1007/s004270000105.

6. Mochizuki K, Nishimiya-Fujisawa C, Fujisawa T. Universal occurrence of the *vasa*-related genes among metazoans and their germline expression in *Hydra*. Dev Genes Evol. 2001;211: 299–308. doi:10.1007/s004270100156.

7. Funayama N. The stem cell system in demosponges: Insights into the origin of somatic stem cells. Dev Growth Differ. 2010;52: 1–14. doi:10.1111/j.1440-169x.2009.01162.x.

8. Bosch TCG, David CN. Stem cells of *Hydra magnipapillata* can differentiate into somatic cells and germ line cells. Dev Biol. 1987;121: 182–191. doi:10.1016/0012-1606(87)90151-5.

9. Fierro-Constaín L, Schenkelaars Q, Gazave E, Haguenauer A, Rocher C, Ereskovsky A, et al. The Conservation of the Germline Multipotency Program, from Sponges to Vertebrates: A Stepping Stone to Understanding the Somatic and Germline Origins. Genome Biol Evol. 2017; evw289. doi:10.1093/gbe/evw289.

10. Müller WA, Teo R, Frank U. Totipotent migratory stem cells in a hydroid. Dev Biol. 2004;275: 215–224. doi:10.1016/j.ydbio.2004.08.006.

11. DuBuc TQ, Schnitzler CE, Chrysostomou E, McMahon ET, Febrimarsa, Gahan JM, et al. Transcription factor AP2 controls cnidarian germ cell induction. Science. 2020;367: 757–762. doi:10.1126/science.aay6782.

12. Baguñà J, Saló E, Auladell C. Regeneration and pattern formation in planarians. III. Evidence that neoblasts are totipotent stem cells and the source of blastema cells. Development. 1989;107: 77–86.

13. Wagner DE, Wang IE, Reddien PW. Clonogenic neoblasts are pluripotent adult stem cells that underlie planarian regeneration. Science. 2011;332: 811–816. doi:10.1126/science.1203983.

14. Newmark PA, Sánchez Alvarado A. Bromodeoxyuridine specifically labels the regenerative stem cells of planarians. Dev Biol. 2000;220: 142–153. doi:10.1006/dbio.2000.9645.

15. Juliano CE, Swartz SZ, Wessel GM. A conserved germline multipotency program. Development. 2010;137: 4113–4126. doi:10.1242/dev.047969.

16. Ewen-Campen B, Schwager EE, Extavour CGM. The molecular machinery of germ line specification. Mol Reprod Dev. 2010;77: 3–18. doi:10.1002/mrd.21091.

17. Nakagawa H, Ishizu H, Hasegawa R, Kobayashi K, Matsumoto M. *Drpiwi-1* is essential for germline cell formation during sexualization of the planarian *Dugesia ryukyuensis*. Dev Biol. 2012;361: 167–176. doi:10.1016/j.ydbio.2011.10.014.

18. Palakodeti D, Smielewska M, Lu Y-C, Yeo GW, Graveley BR. The PIWI proteins SMEDWI-2 and SMEDWI-3 are required for stem cell function and piRNA expression in planarians. RNA. 2008;14: 1174–1186. doi:10.1261/rna.1085008.

19. Reddien PW, Oviedo NJ, Jennings JR, Jenkin JC, Sánchez Alvarado A. SMEDWI-2 is a PIWI-like protein that regulates planarian stem cells. Science. 2005;310: 1327–1330. doi:10.1126/science.1116110.

20. Iyer H, Issigonis M, Sharma PP, Extavour CG, Newmark PA. A premeiotic function for *boule* in the planarian *Schmidtea mediterranea*. Proc Natl Acad Sci USA. 2016;113: E3509–18. doi:10.1073/pnas.1521341113.

21. Salvetti A, Rossi L, Lena A, Batistoni R, Deri P, Rainaldi G, et al. *DjPum*, a homologue of *Drosophila Pumilio*, is essential to planarian stem cell maintenance. Development. 2005;132: 1863–1874. doi:10.1242/dev.01785.

22. Solana J, Lasko P, Romero R. *Spoltud-1* is a chromatoid body component required for planarian long-term stem cell self-renewal. Dev Biol. 2009;328: 410–421. doi:10.1016/j.ydbio.2009.01.043.

23. Rouhana L, Shibata N, Nishimura O, Agata K. Different requirements for conserved post-transcriptional regulators in planarian regeneration and stem cell maintenance. Dev Biol. 2010;341: 429–443. doi:10.1016/j.ydbio.2010.02.037.

24. Shibata N, Kashima M, Ishiko T, Nishimura O, Rouhana L, Misaki K, et al. Inheritance of a Nuclear PIWI from Pluripotent Stem Cells by Somatic Descendants Ensures Differentiation by Silencing Transposons in Planarian. Dev Cell. 2016;37: 226–237. doi:10.1016/j.devcel.2016.04.009.

25. Wagner DE, Ho JJ, Reddien PW. Genetic regulators of a pluripotent adult stem cell system in planarians identified by RNAi and clonal analysis. Cell Stem Cell. 2012;10: 299–311. doi:10.1016/j.stem.2012.01.016.

26. Guo T, Peters AHFM, Newmark PA. A *bruno*-like gene is required for stem cell maintenance in planarians. Dev Cell. 2006;11: 159–169. doi:10.1016/j.devcel.2006.06.004.

27. Kim IV, Duncan EM, Ross EJ, Gorbovytska V, Nowotarski SH, Elliott SA, et al. Planarians recruit piRNAs for mRNA turnover in adult stem cells. Genes Dev. 2019;33: 1575–1590. doi:10.1101/gad.322776.118.

28. Issigonis M, Newmark PA. From worm to germ: Germ cell development and regeneration in planarians. Curr Top Dev Biol. 2019;135: 127–153. doi:10.1016/bs.ctdb.2019.04.001.

29. Newmark PA, Wang Y, Chong T. Germ cell specification and regeneration in planarians. Cold Spring Harbor symposia on quantitative biology. 2008;73: 573–581. doi:10.1101/sqb.2008.73.022.

30. Wolff E. Recent researches on the regeneration of Planaria. In: Rudnick D, editor. Regeneration: 20th Growth Symposium, Vol. 20. New York: The Roland Press; 1962. pp. 53– 84.

31. Wang J, Chen R, Collins JJ. Systematically improved in vitro culture conditions reveal new insights into the reproductive biology of the human parasite *Schistosoma mansoni*. PLOS Biol. 2019;17: e3000254. doi:10.1371/journal.pbio.3000254.

32. Wang J, Collins JJ. Identification of new markers for the *Schistosoma m*ansoni vitelline lineage. Int J Parasitol. 2016;46: 405–410. doi:10.1016/j.ijpara.2016.03.004.

33. Takahashi K, Yamanaka S. Induction of pluripotent stem cells from mouse embryonic and adult fibroblast cultures by defined factors. Cell. 2006;126: 663–676. doi:10.1016/j.cell.2006.07.024.

34. Wang Y, Zayas RM, Guo T, Newmark PA. *nanos* function is essential for development and regeneration of planarian germ cells. Proc Natl Acad Sci USA. 2007;104: 5901–5906. doi:10.1073/pnas.0609708104.

35. Chong T, Collins JJ, Brubacher JL, Zarkower D, Newmark PA. A sex-specific transcription factor controls male identity in a simultaneous hermaphrodite. Nat Comm. 2013;4: 1814. doi:10.1038/ncomms2811.

36. Handberg-Thorsager M, Saló E. The planarian *nanos*-like gene *Smednos* is expressed in germline and eye precursor cells during development and regeneration. Dev Genes and Evol. 2007;217: 403–411. doi:10.1007/s00427-007-0146-3.

37. Sato K, Shibata N, Orii H, Amikura R, Sakurai T, Agata K, et al. Identification and origin of the germline stem cells as revealed by the expression of *nanos*-related gene in planarians. Develop Growth Differ. 2006;48: 615–628. doi:10.1111/j.1440-169x.2006.00897.x.

38. Farnesi RM, Marinelli M, Tei S, Vagnetti D. Ultrastructural research on the spermatogenesis in *Dugesia lugubris*. SL Riv Di Biol. 1977;70: 113–36.

39. Saberi A, Jamal A, Beets I, Schoofs L, Newmark PA. GPCRs direct germline development and somatic gonad function in planarians. PLOS Biol. 2016;14: e1002457. doi:10.1371/journal.pbio.1002457.

40. Rouhana L, Tasaki J, Saberi A, Newmark PA. Genetic dissection of the planarian reproductive system through characterization of *Schmidtea mediterranea* CPEB homologs. Dev Biol. 2017;426: 43–55. doi:10.1016/j.ydbio.2017.04.008.

41. Morgan TH. Growth and regeneration in *Planaria lugubris*. Arch Entwickl Org. 1902;13: 179–212. doi:10.1007/bf02161982.

42. Chong T, Stary JM, Wang Y, Newmark PA. Molecular markers to characterize the hermaphroditic reproductive system of the planarian *Schmidtea mediterranea*. BMC Dev Biol. 2011;11: 69. doi:10.1186/1471-213x-11-69.

43. Davies EL, Lei K, Seidel CW, Kroesen AE, McKinney SA, Guo L, et al. Embryonic origin of adult stem cells required for tissue homeostasis and regeneration. eLife. 2017;6. doi:10.7554/elife.21052.

44. Hyman LH. The Invertebrates: Vol. 2, Platyhelminthes and Rhynchocoela, the Acoelomate Bilateria. McGraw-Hill; 1951.

45. King RS, Newmark PA. In situ hybridization protocol for enhanced detection of gene expression in the planarian *Schmidtea mediterranea*. BMC Dev Biol. 2013;13: 8–16. doi:10.1186/1471-213x-13-8.

46. King RS, Newmark PA. Whole-mount in situ hybridization of planarians. Methods Mol Biol. (Clifton, NJ). 2018;1774: 379–392. doi:10.1007/978-1-4939-7802-1_12.

47. Laumer CE, Giribet G. Inclusive taxon sampling suggests a single, stepwise origin of ectolecithality in Platyhelminthes. Biol J Linn Soc. 2014;111: 570–588. doi:10.1111/bij.12236.

48. Laumer CE, Hejnol A, Giribet G. Nuclear genomic signals of the ‘microturbellarian’ roots of platyhelminth evolutionary innovation. eLife. 2015;4: e05503. doi:10.7554/elife.05503.

49. Egger B, Lapraz F, Tomiczek B, Müller S, Dessimoz C, Girstmair J, et al. A transcriptomic-phylogenomic analysis of the evolutionary relationships of flatworms. Curr Biol. 2015;25: 1347– 1353. doi:10.1016/j.cub.2015.03.034.

50. Wang Y, Stary JM, Wilhelm JE, Newmark PA. A functional genomic screen in planarians identifies novel regulators of germ cell development. Genes Dev. 2010;24: 2081–2092. doi:10.1101/gad.1951010.

51. Collins JJ, Hou X, Romanova EV, Lambrus BG, Miller CM, Saberi A, et al. Genome-wide analyses reveal a role for peptide hormones in planarian germline development. PLOS Biol. 2010;8: e1000509. doi:10.1371/journal.pbio.1000509.

52. Zayas RM, Hernández A, Habermann B, Wang Y, Stary JM, Newmark PA. The planarian *Schmidtea mediterranea* as a model for epigenetic germ cell specification: analysis of ESTs from the hermaphroditic strain. Proc Natl Acad Sci USA. 2005;102: 18491–18496. doi:10.1073/pnas.0509507102.

53. Rouhana L, Vieira AP, Roberts-Galbraith RH, Newmark PA. PRMT5 and the role of symmetrical dimethylarginine in chromatoid bodies of planarian stem cells. Development. 2012;139: 1083–1094. doi:10.1242/dev.076182.

54. Hayashi K, Kobayashi T, Umino T, Goitsuka R, Matsui Y, Kitamura D. SMAD1 signaling is critical for initial commitment of germ cell lineage from mouse epiblast. Mech Dev. 2002;118: 99–109. doi: 10.1016/s0925-4773(02)00237-x.

55. Lawson KA, Dunn NR, Roelen BA, Zeinstra LM, Davis AM, Wright CV, et al. *Bmp4* is required for the generation of primordial germ cells in the mouse embryo. Genes Dev. 1999;13: 424–436. doi: 10.1101/gad.13.4.424.

56. Ying Y, Liu XM, Marble A, Lawson KA, Zhao GQ. Requirement of *Bmp8b* for the generation of primordial germ cells in the mouse. Mol. Endocrinol. 2000;14: 1053–1063. doi:10.1210/mend.14.7.0479.

57. Ying Y, Zhao G-Q. Cooperation of endoderm-derived BMP2 and extraembryonic ectoderm-derived BMP4 in primordial germ cell generation in the mouse. Dev Biol. 2001;232: 484–492. doi:10.1006/dbio.2001.0173.

58. Chang DH, Cattoretti G, Calame KL. The dynamic expression pattern of B lymphocyte induced maturation protein-1 (Blimp-1) during mouse embryonic development. Mech Dev. 2002;117: 305–309. doi:10.1016/s0925-4773(02)00189-2.

59. Hayashi K, Lopes SMC de S, Surani MA. Germ cell specification in mice. Science. 2007;316: 394–396. doi:10.1126/science.1137545.

60. Ohinata Y, Payer B, O’Carroll D, Ancelin K, Ono Y, Sano M, et al. Blimp1 is a critical determinant of the germ cell lineage in mice. Nature. 2005;436: 207–213. doi:10.1038/nature03813.

61. Vincent SD, Dunn NR, Sciammas R, Shapiro-Shalef M, Davis MM, Calame K, et al. The zinc finger transcriptional repressor Blimp1/Prdm1 is dispensable for early axis formation but is required for specification of primordial germ cells in the mouse. Development. 2005;132: 1315– 1325. doi:10.1242/dev.01711.

62. Magnúsdóttir E, Surani MA. How to make a primordial germ cell. Development. 2014;141: 245–252. doi:10.1242/dev.098269.

63. Grabole N, Tischler J, Hackett JA, Kim S, Tang F, Leitch HG, et al. Prdm14 promotes germline fate and naive pluripotency by repressing FGF signalling and DNA methylation. EMBO Rep. 2013;14: 629–637. doi:10.1038/embor.2013.67.

64. Kurimoto K, Yabuta Y, Ohinata Y, Shigeta M, Yamanaka K, Saitou M. Complex genome-wide transcription dynamics orchestrated by Blimp1 for the specification of the germ cell lineage in mice. Genes Dev. 2008;22: 1617–1635. doi:10.1101/gad.1649908.

65. Yamaji M, Seki Y, Kurimoto K, Yabuta Y, Yuasa M, Shigeta M, et al. Critical function of *Prdm14* for the establishment of the germ cell lineage in mice. Nat Genet. 2008;40: 1016–1022. doi:10.1038/ng.186.

66. Magnúsdóttir E, Dietmann S, Murakami K, Günesdogan U, Tang F, Bao S, et al. A tripartite transcription factor network regulates primordial germ cell specification in mice. Nat Cell Biol. 2013;15: 1–13. doi:10.1038/ncb2798.

67. Weber S, Eckert D, Nettersheim D, Gillis AJM, Schäfer S, Kuckenberg P, et al. Critical Function of AP-2gamma/TCFAP2C in Mouse Embryonic Germ Cell Maintenance. Biol Reprod. 2010;82: 214–223. doi:10.1095/biolreprod.109.078717.

68. Nakaki F, Hayashi K, Ohta H, Kurimoto K, Yabuta Y, Saitou M. Induction of mouse germ-cell fate by transcription factors in vitro. Nature. 2013;501: 222–226. doi:10.1038/nature12417.

69. Magnúsdóttir E, Gillich A, Grabole N, Surani MA. Combinatorial control of cell fate and reprogramming in the mammalian germline. Curr Opin Genet Dev. 2012;22: 466–474. doi:10.1016/j.gde.2012.06.002.

70. Nakamura T, Extavour CG. The transcriptional repressor Blimp-1 acts downstream of BMP signaling to generate primordial germ cells in the cricket *Gryllus bimaculatus*. Development. 2016;143: 255–263. doi:10.1242/dev.127563.

71. Donoughe S, Nakamura T, Ewen-Campen B, Green DA, Henderson L, Extavour CG. BMP signaling is required for the generation of primordial germ cells in an insect. Proc Natl Acad Sci USA. 2014;111: 4133–4138. doi:10.1073/pnas.1400525111.

72. Yabuta Y, Kurimoto K, Ohinata Y, Seki Y, Saitou M. Gene expression dynamics during germline specification in mice identified by quantitative single-cell gene expression profiling. Biol Reprod. 2006;75: 705–716. doi:10.1095/biolreprod.106.053686.

73. Kitadate Y, Shigenobu S, Arita K, Kobayashi S. Boss/Sev signaling from germline to soma restricts germline-stem-cell-niche formation in the anterior region of *Drosophila* male gonads. Dev Cell. 2007;13: 151–159. doi:10.1016/j.devcel.2007.05.001.

74. Kitadate Y, Kobayashi S. Notch and Egfr signaling act antagonistically to regulate germ-line stem cell niche formation in *Drosophila* male embryonic gonads. Proc Natl Acad Sci USA. 2010;107: 14241–14246. doi:10.1073/pnas.1003462107.

75. Gilboa L, Lehmann R. Soma–germline interactions coordinate homeostasis and growth in the *Drosophila* gonad. Nature. 2006;443: 97–100. doi:10.1038/nature05068.

76. Martín-Durán JM, Egger B. Developmental diversity in free-living flatworms. Evodevo. 2012;3: 7. doi:10.1186/2041-9139-3-7.

77. Cebrià F, Newmark PA. Planarian homologs of netrin and netrin receptor are required for proper regeneration of the central nervous system and the maintenance of nervous system architecture. Development. 2005;132: 3691–3703. doi:10.1242/dev.01941.

78. Sánchez Alvarado A, Newmark PA, Robb SM, Juste R. The *Schmidtea mediterranea* database as a molecular resource for studying platyhelminthes, stem cells and regeneration. Development. 2002;129: 5659–5665. doi: 10.1242/dev.00167.

79. Rouhana L, Weiss JA, Forsthoefel DJ, Lee H, King RS, Inoue T, et al. RNA interference by feeding in vitro-synthesized double-stranded RNA to planarians: methodology and dynamics. Dev Dyn. 2013;242: 718–730. doi:10.1002/dvdy.23950.

